# An amortized approach to non-linear mixed-effects modeling based on neural posterior estimation

**DOI:** 10.1101/2023.08.22.554273

**Authors:** Jonas Arruda, Yannik Schälte, Clemens Peiter, Olga Teplytska, Ulrich Jaehde, Jan Hasenauer

## Abstract

Non-linear mixed-effects models are a powerful tool for studying heterogeneous populations in various fields, including biology, medicine, economics, and engineering. Here, the aim is to find a distribution over the parameters that describe the whole population using a model that can generate simulations for an individual of that population. However, fitting these distributions to data is computationally challenging if the description of individuals is complex and the population is large. To address this issue, we propose a novel machine learning-based approach: We exploit neural density estimation based on conditional normalizing flows to approximate individual-specific posterior distributions in an amortized fashion, thereby allowing for efficient inference of population parameters. Applying this approach to problems from cell biology and pharmacology, we demonstrate its unseen flexibility and scalability to large data sets compared to established methods.

## 1. Introduction

Heterogeneity within populations is a common phenomenon in various fields, including epidemiology, pharmacology, ecology, and economics. It is, for instance, well-established that the human immune system exhibits substantial variability among individuals (Liston et al., 2021; Brodin & Davis, 2017), that individual patients respond differently to treatments (Claret et al., 2009; Ribba et al., 2014; Groenland et al., 2019), that genetically identical cells develop pronounced cell-to-cell variability (Spencer et al., 2009; Swain et al., 2002), but also that individual students show a broad spectrum of abilities (Goldstein, 1987). This heterogeneity can be described and analyzed using *non-linear mixed-effects (NLME)* models, a powerful class of statistical tools. NLME models can account for similarities and differences between individuals using fixed effects, random effects, and covariates. This allows for a high degree of flexibility and interpretability. These models are widely used for statistical analysis (Yu et al., 2022; Llamosi et al., 2016), hypothesis testing (Bortz & Nelson, 2006), and predictions (Claret et al., 2009; Ribba et al., 2014).

NLME models depend on unknown parameters, such as reaction rates and initial concentrations, which often need to be estimated from data. Estimating these parameters – often also called *parameter inference* – provides key insights about the data and the underlying processes. The main challenge in inferring these parameters lies in the likelihood formulation at the individual level. For this, there is generally no closed-form solution (Pinheiro, 1994). Particularly for large populations, this becomes a problem, as the required marginalization must be performed for all individuals.

Here, we present a new approach based on invertible neural networks to estimate the parameters of NLME models. We use simulation-based neural posterior estimation, which has been developed to address general parameter estimation problems (Cranmer et al., 2020). We train a mapping – a conditional normalizing flow – from a latent distribution to individual-specific posteriors conditioned on observed individual-level data. During the training of this neural posterior estimator, only simulations of a generative model and no real data are used. In the following inference phase, the trained estimator can be applied highly efficiently to any similar data set with different distributions of individuals in the population without any further simulations, facilitating the estimation of NLME model parameters in an amortized fashion. On problems from cell biology and pharmacology, we compare our method with state-of-the-art and widely used techniques in the field of NLME models: the stochastic approximation expectation maximization algorithm (SAEM) (Kuhn & Lavielle, 2005) implemented in `Monolix` (Lixoft SAS, 2023) and the first-order conditional estimation with interaction (FOCEI) (Wang, 2008) implemented in `NONMEM` (Beal & Sheiner, 1980).

## 2. Method

Our novel approach to parameter inference for non-linear mixed-effects (NLME) models consists of three phases (summarized in Figure 1):

**Figure 1.**
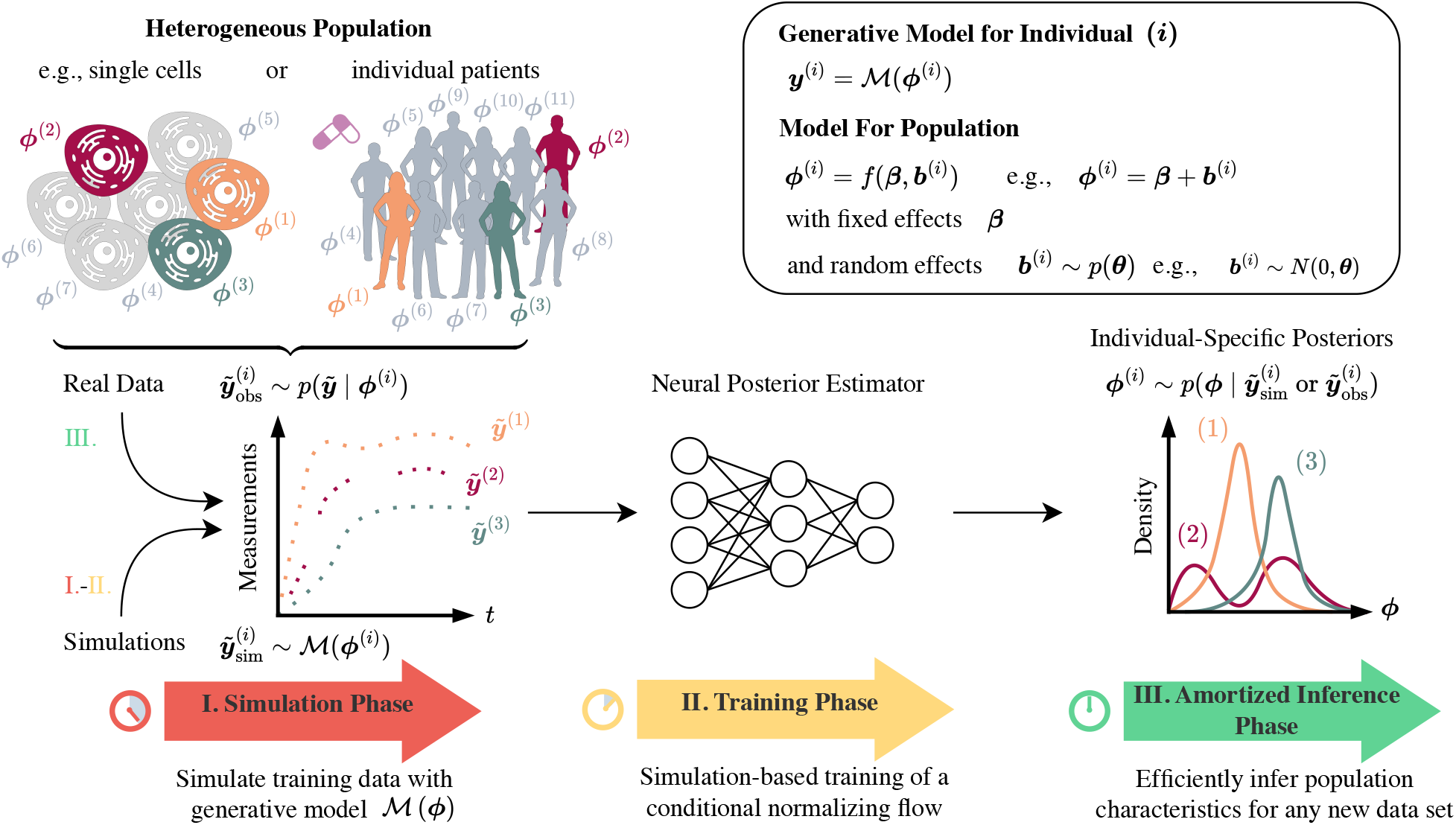
Three phases of the amortized approach. (**I**.) The simulation phase, where we generate data from the model ℳ(*ϕ*), (**II**.) the training phase, where we train the neural posterior estimator to predict individual-specific posteriors based on the simulations, and (**III**.) the amortized inference phase, where we infer the population parameters of the non-linear mixed-effects model given observed data.

I. In the simulation phase, samples from a prior are generated to produce a set of simulations and individual-specific parameters using a generative model. Each simulation belongs to a synthetic individual in a population. Simulations can also be performed online during the next phase.
II. In the training phase, a global approximation of the posterior distribution is learned for any pair of simulations and parameters. This neural posterior estimator can then predict individual-specific posteriors.
III. In the amortized inference phase, a population model is assumed and population-level characteristics are inferred using an efficient approximation to the population likelihood based on samples of every individual in the population from the neural posterior estimator.

### 2.1. Basic Definitions

We consider a population of individuals *i* ∈ *{*1, …, *N}, N* ∈ ℕ, for example, a group of people or an ensemble of single cells. For each of them, we have *n* ∈ ℕ noisy measurements 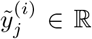 at time points *t*_*j*_ ∈ ℝ_*≥*0_ for *j* ∈ {1, …, *n*} . To account for errors introduced during the measurement process, a noise model with i.i.d. measurement noise is assumed. In the simplest case this is 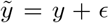, where *ϵ* ∼ *𝒩* (0, *σ*^2^) is the measurement error with variance *σ*^2^. Then, our observed data is the set 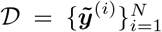 . We can extend this framework to account for multiple or censored measurements at each time point and different time points for each individual.

### The generative model

ℳ (***ϕ***) can generate simulation 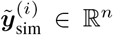 for a given set of parameters ***ϕ*** ∈ Ω ⊆ ℝ^*k*^ and time points ***t***. As a generative model, we understand any parametric model, such as linear models, the solution of (stochastic) differential equations, or Markov jump processes, which can produce simulations for an individual *i* given some parameters ***ϕ*** and time points ***t***. In our case, this will be the (numerical) solution of a differential equation, and we assume that the noise model is part of ℳ.

### The non-linear mixed-effects (NLME) model

is a popular way to describe observations of the entire population using the generative model ℳ and individual-specific parameters ***ϕ***^(*i*)^ ∈ Ω⊆ *ℝ*^*k*^. Therefore, in NLME models, it is assumed that the population can be described by unknown fixed effects ***β***, and the distribution of unknown random effects ***b***^(*i*)^ specific to each individual *i* (Pinheiro, 1994).

Commonly, fixed and random effects are linked as a lin-ear combination to the individual-specific parameters ***ϕ***^(*i*)^. Here, we link these effects to individual-specific parameters using a *population model f*, such that ***ϕ***^(*i*)^ = *f* (***β, b***^(*i*)^), where the function *f* can be any function injective in each variable. It is possible to extend the population model to include covariates, that is, additional observed information for each individual *i*.

For ease of notation, we consider a single vector of population parameters ***θ*** that fully characterize the distribution of individual-specific parameters ***ϕ***^(*i*)^. For example, we can represent ***θ*** as a pair of fixed effects ***β*** and the covariance matrix **Ψ** of random effects if ***b***^(*i*)^ ∼ 𝒩 (0, **Ψ**) and ***ϕ***^(*i*)^ = *f* (***β, b***^(*i*)^). In general, we allow the random effects to follow any probability distribution that admits a density, such as (log)-normal distributions, Cauchy distributions, or mixture distributions.

### 2.2. Joint Likelihood

Our objective is to find the optimal population parameters ***θ***^*^ that maximize the joint likelihood *p*(𝒟|***θ***) of all individuals in the population, which is

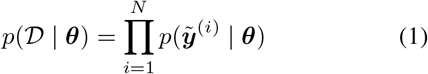

since we assumed that measurement noise is i.i.d.

Based on the population model and the distribution of random effects, we induce a population distribution *p*_pop_(***ϕ*** | ***θ***). For a linear population model ***ϕ***^(*i*)^ = ***β*** + ***b***^(*i*)^ and ***b***^(*i*)^ ∼ 𝒩 (0, **Ψ**), *p*_pop_ is the density of a normal random variable with mean **β** and covariance matrix **Ψ**. Using the population distribution, we can thus express the likelihood for every individual *i* by the law of total probability as

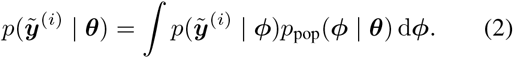

The marginal likelihood 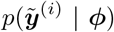 is implicitly induced by ℳ including the respective noise model. Usually, the likelihood is not fully tractable if the generative model is stochastic.

Since ***ϕ*** is unknown, we need to marginalize the individual-specific parameters out and solve the integral in (2) for every individual separately. In general, this integral has no closed-form solution, hence solving this marginalization efficiently is the main challenge in parameter inference in non-linear mixed-effects models (Pinheiro, 1994).

### 2.3. Individual-specific neural posterior estimator

In the following, we develop an approach to efficiently maximize *p*(𝒟| ***θ***) if we can sample from the individual specific posterior distributions 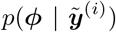. In general, individual measurements are not sufficiently informative to obtain reliable population estimates, and only joint information is reliable (Pinheiro, 1994). However, using a Bayesian approach to describe individuals, we encode all available information on a specific individual *i* in the posterior of the parameters ***ϕ***^(*i*)^ and then combine samples from this posterior to infer the characteristics of the population. Following this dogma of Bayesian statistics, all parameters – also those which are considered fixed across the population – will first be treated as random variables.

For that, let *p*(***ϕ***) be a prior density that encodes prior knowledge for individual specific parameters. Then, using the product rule, we get the relation of likelihood and posterior density 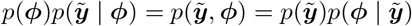.

### Conditional normalizing flows

can transform a complicated conditional density, such as a posterior probability, into a simpler density from which we know how to sample. This method allows for efficient and accurate sampling and density evaluation (Rezende & Mohamed, 2015; Papamakarios et al., 2021).

We introduce latent variables ***z*** described by a multivariate normal distribution *p*(***z***). The parameters ***ϕ*** are mapped to these latent variables conditional on measurements 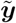 by a conditional normalizing flow 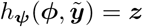 . This invertible transformation is parameterized by ***ψ*** and has a tractable Jacobian by construction. The approximation *q*_***ψ***_ to the target density 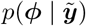 is given by the change-of-variables formula

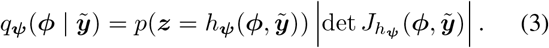

If we know *h*_***ψ***_, we can sample from1 the posterior 11by sampling ***z*** ∼ *p*(***z***) and applying 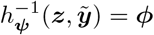. We call *q*_*ψ*_ a *neural posterior estimator*.

### To train the conditional normalizing flow

*h*_***ψ***_, we minimize the Kullback-Leibler divergence between the true and approximate posterior distributions as in (Papamakarios et al., 2017; Le et al., 2017):

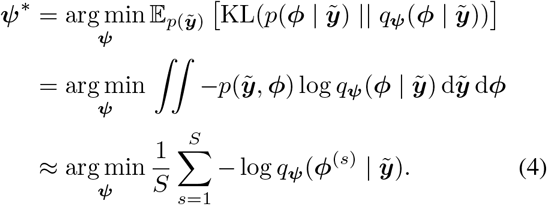

To estimate the integral, we need samples from the joint distribution 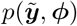. Sampling from the prior distribution ***ϕ*** ∼ *p*(***ϕ***) and simulating using ℳ(***ϕ***) corresponds to sampling from this joint density. Using the transformation in (3), the approximation in (4) can be efficiently evaluated.

#### Algorithm 1 Amortized Inference for NLME Models

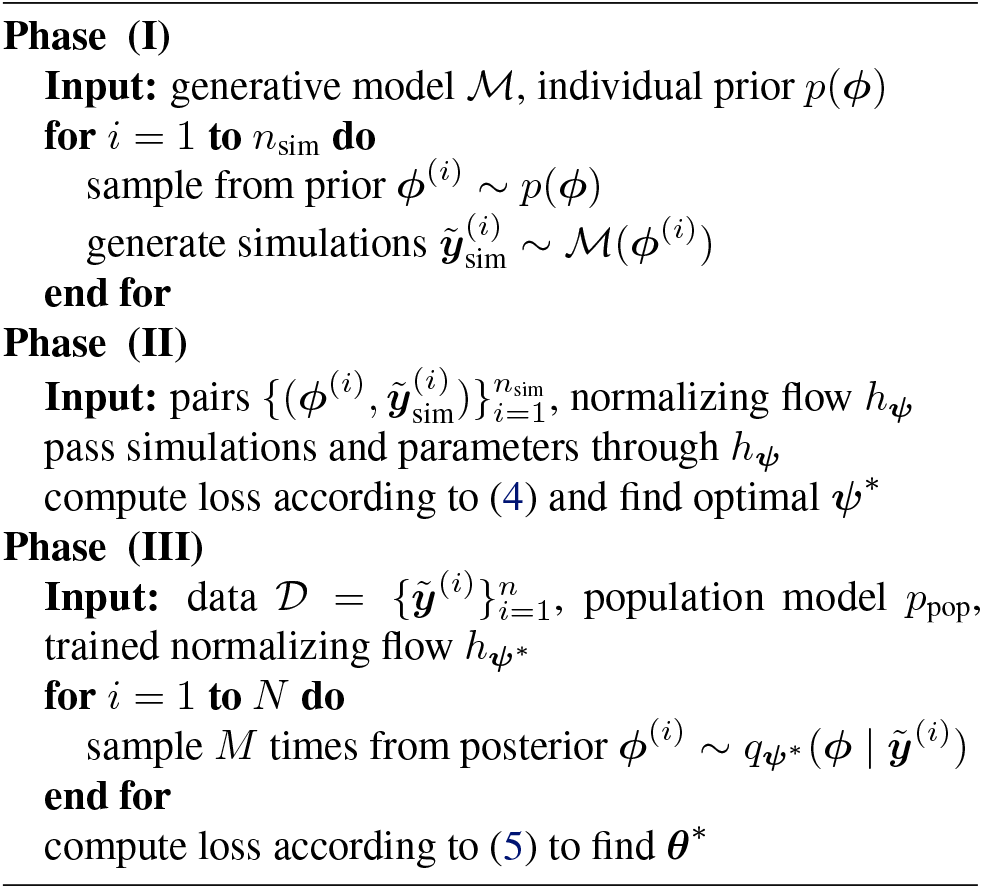

By minimizing (4), we train a global approximation of the posterior distribution 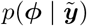 for any parameters and data 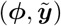. In particular, we parameterize the conditional normalizing flow by an invertible neural network using neural spline flows (Durkan et al., 2019) and learned summary statistics of 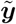.

### Summary statistics

are a low-dimensional representation of 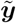 and learned by a flexible summary network that is trained together with the conditional normalizing flow. In (Papamakarios et al., 2017; Le et al., 2017), it was shown that the samples transformed backward from *p*(***z***) will follow the true posterior if the conditional normalizing flow and the summary network are expressive enough. In particular, we use long short-term memory neural networks (LSTMs) for time trajectories (Hochreiter & Schmidhuber, 1997). This ensures that regardless of the number of observations, we get a fixed length vector of summary statistics.

### 2.4. Problem reformulation allows use of posterior density

We note that the likelihood in (2) can be written as a conditional expectation over individual-specific posteriors

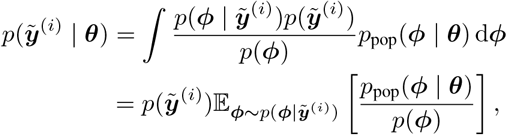

given that the prior *p*(***ϕ***) is non-zero in the support of ***ϕ***. If we can sample from the posterior distribution 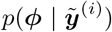, we can construct a Monte Carlo estimator for the likelihood

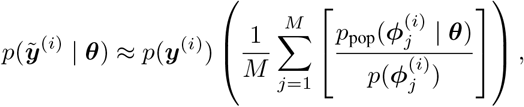

with 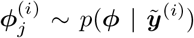 i.i.d. for *j* = 1, …, *M* for each individual *i*.

Taking the logarithm of (1), we can drop the additive term *p*(***y***) which is independent of ***θ*** and find the optimal population parameters ***θ***^*^ by solving the maximization problem

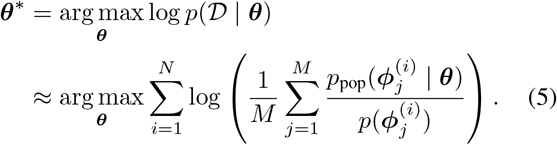

In general, the Monte Carlo approximation to an integral is unbiased, and the error rate of the approximation 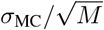 depends on the sample size *M* and the variance 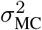 of the ratio *p*_pop_(***ϕ*** |***θ***)*/p*(***ϕ***) (Robert & Casella, 2004). From this, we directly get the following proposition.

#### Proposition 2.1.

*Assume that* ***ϕ***^(*i*)^ ∼ *p*_*pop*_(***ϕ*** | ***θ***), *that the prior p*(***ϕ***) *is non-zero in the support of* ***ϕ*** *and that we can sample from the true posteriors* 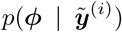 *for every individual i. Then*, ***θ*** *converges to the true maximum likelihood estimate* ***θ***^*^ *as the sample size M → ∞*.

The variance 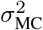 depends only on the ratio of *p*_pop_(***ϕ***|***θ***) and *p*(***ϕ***). Therefore, the prior has the role of an importance weight and should be selected to have a shape similar to the population distribution. This decreases the number of samples *M* we need to get a sufficiently good approximation of the likelihood (see Supplement A.4.1 for further discussion).

The maximization problem (5) can be solved by numerical optimization using samples from the neural posterior esti-Mator 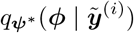. The optimization is computationally efficient and simple, as no numerical simulations of the underlying model are required. Hence, the computational costs of inferring population parameters are negligible.

We show our novel three-phase procedure for the inference of NLME models in Algorithm 1. By repeating phase (III) using different population models or data sets, we amortize the computational cost of phases (I) and (II). Repeated inference can be desirable due to hypothesis testing of various population models or repeated experiments.

## 3. Results

The proposed approach to fitting the NLME models is based on the approximation of the individual-specific posterior distributions with conditional normalizing flows. As the accuracy of these approximations is critical, we assessed in a first step the approximation quality (see Supplement A.3). We considered two published ODE-based NLME models of mRNA transfection with measured single-cell data (Fröhlich et al., 2018). These ODE models describe the transfection process (Figure 2A) – which is at the core of modern mRNA vaccines (Pardi et al., 2018) – at the single-cell level. The solution to these ODEs corresponds to our generative model ℳ. The models possess, respectively, six and eleven parameters ***ϕ*** that describe two and four hidden state variables (Figure 2B, see Supplement A.1 for details on the models). The measurements consist of dense temporally resolved fluorescence intensities of different single-cells, which were transfected with mRNA coding for a green fluorescent protein (GFP), and measured for 30 hours using micropatterned protein arrays and time-lapse microscopy (Figure 2C). For each ODE model, we simulate data and train a neural posterior estimator accordingly with phase (I) and (II) from our proposed Algorithm 1. Subsequently, we examine a second application from pharmacokinetics, which is a major application area for NLME models.

**Figure 2.**
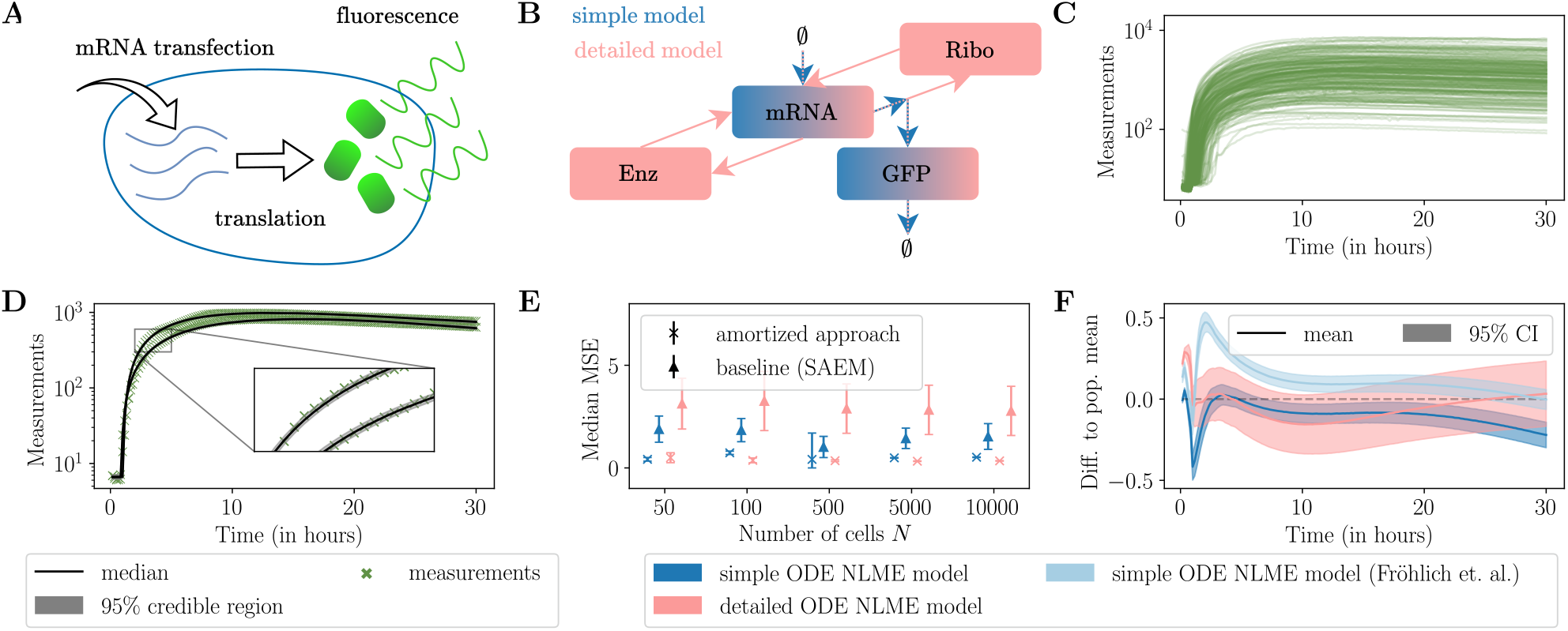
Validation of the amortized approach on single-cell NLME models. (**A**) Schematic representation of GFP translation after mRNA transfection in a single cell. (**B**) Visualization of the simple and detailed single-cell ODE models, where the color indicates the states included in the respective model (see Supplement A.1 for details on the models). (**C**) Fluorescence intensity time courses of 200 single cells out of 5488. (**D**) Credible regions of trajectories (simple single-cell ODE model) estimated by the neural posterior estimator for two real cells. (**E**) Median of the mean squared error (MSE) of the estimated compared to the true parameters of the synthetic data for both single-cell NLME models is shown for different numbers of cells (*M* = 100 posterior samples used per cell, median of 100 runs *±* one standard deviation). (**F**) The difference in the population mean estimated from real trajectories and simulations generated with the estimated population parameters is shown with a 95% confidence interval (CI). In addition to the models fitted with the amortized approach, the best fit of Fröhlich et al. (2018) for the simple ODE model is shown (Fröhlich et al., 2018).

We consider two scenarios to validate our method using the same trained neural posterior estimator: a) using synthetic data where we want to recover the true sample parameters, and b) using real data 𝒟 = 𝒟 _eGFP_ ∪ 𝒟 _d2eGFP_, with two distinct variants of GFP, namely eGFP and d2eGFP, that differ in their protein lifetime (Fröhlich et al., 2018). For the synthetic data, we assume a log-normal distribution with diagonal covariance matrix **Ψ** for all parameters, that is, ***ϕ***∼ log 𝒩 (***β*, Ψ**) and ***θ*** = (***β*, Ψ**). For the real data and both variants of GFP, we can use the same generative model ℳ since we assume that they only differ in the parameter that describes GFP degeneration (indexed by *γ*). We encode the variant as a binary covariate *c*^(*i*)^ ∈ {0, 1}. Hence, we have ***ϕ***^(*i*)^ ∼ log 𝒩 (***β*, Ψ**) *c*^(*i*)^, where the elements ***β***^*γ*^ and **Ψ**_*γγ*_ depend on the respective variant of the single-cell *i*. Then, for each element 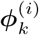 the population model is

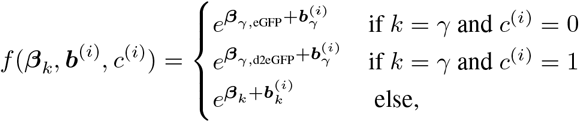

where ***b***^(*i*)^ ∼ 𝒩 (0, **Ψ**) | *c*^(*i*)^.

We implemented conditional normalizing flows using the `BayesFlow` tool (Radev et al., 2023). We assessed the quality of the neural posteriors by comparing them to approximations using MCMC. For synthetic data, we got nearly ideal agreement while on real data agreement up to model misspecification (see Supplement A.3). To estimate population parameters, we implemented the optimization problem (5) as an objective function in the `pyPESTO` tool-box (Schälte et al., 2023) and used the local optimization method L-BFGS (Liu & Nocedal, 1989) implemented in `SciPy` (Virtanen et al., 2020). For further details on the neural network architecture, we refer to the Supplement A.3. We compare our method to the `Monolix` (Lixoft SAS, 2023) implementation of the state-of-the-art method SAEM, which is an unbiased algorithm and converges under very general conditions (Kuhn & Lavielle, 2005).

### 3.1. Machine learning-based approach provides accurate estimates of population parameters

Given the accurate approximation of posteriors on an individual-specific level (Figure 2D and Supplement A.3), we can use the pre-trained densities to estimate the NLME population parameters ***θ***. To assess the accuracy of our approach, we generated synthetic data using the two NLME models of mRNA transfection (see Supplement A.1.1), and compared the mean squared distance of the true parameters to the estimated parameters of our approach and to the estimated parameters of the state-of-the-art method SAEM (Kuhn & Lavielle, 2005). Moreover, we compared our results with those published in (Fröhlich et al., 2018), which used a Laplacian approximation to the population likelihood to reduce the computational costs compared to SAEM.

Our comparisons show that, for different data set sizes and models, our method was able to recover the true parameters with a lower recovery error than SAEM and a smaller variance in the estimates (Figure 2E). For each ODE model, we trained only one neural posterior estimator, which could be used for inference on all different single-cell data sets, while SAEM needed a full restart for each data set. In addition, the estimated population means for both models of our amortized approach show trajectories similar to those published in (Fröhlich et al., 2018) for the real data (Figure 2F), with only a minor shift observed in the case of the simple model. Furthermore, we can confirm the result of (Fröhlich et al., 2018) that the detailed model describes the initial fluorescence activity more accurately (Supplement A.4).

In summary, our approach based on amortized neural posterior estimation was able to provide accurate estimates of population parameters for synthetic and real data, while we used the same neural posterior estimator for each model.

### 3.2. Amortization for large populations, new data sets and changing population models achieved

Data simulation and training of the neutral posterior estimator are the most computationally demanding phases. For both phases, the detailed NLME model required twice as much computation time compared to the simple model. However, the subsequent inference phase of population parameters is highly efficient for a new data set.

Our method requires two orders of magnitude less computation time compared to SAEM and has a slower increase with respect to the number of individuals in the population (Figure 3A). In particular, if the population was large (10, 000 cells in this case), we already amortized the training time cost compared to SAEM for a single data set. In total, for all data sets and models together, SAEM had an overall computation time 112 times longer than our amortized approach including all three phases.

**Figure 3.**
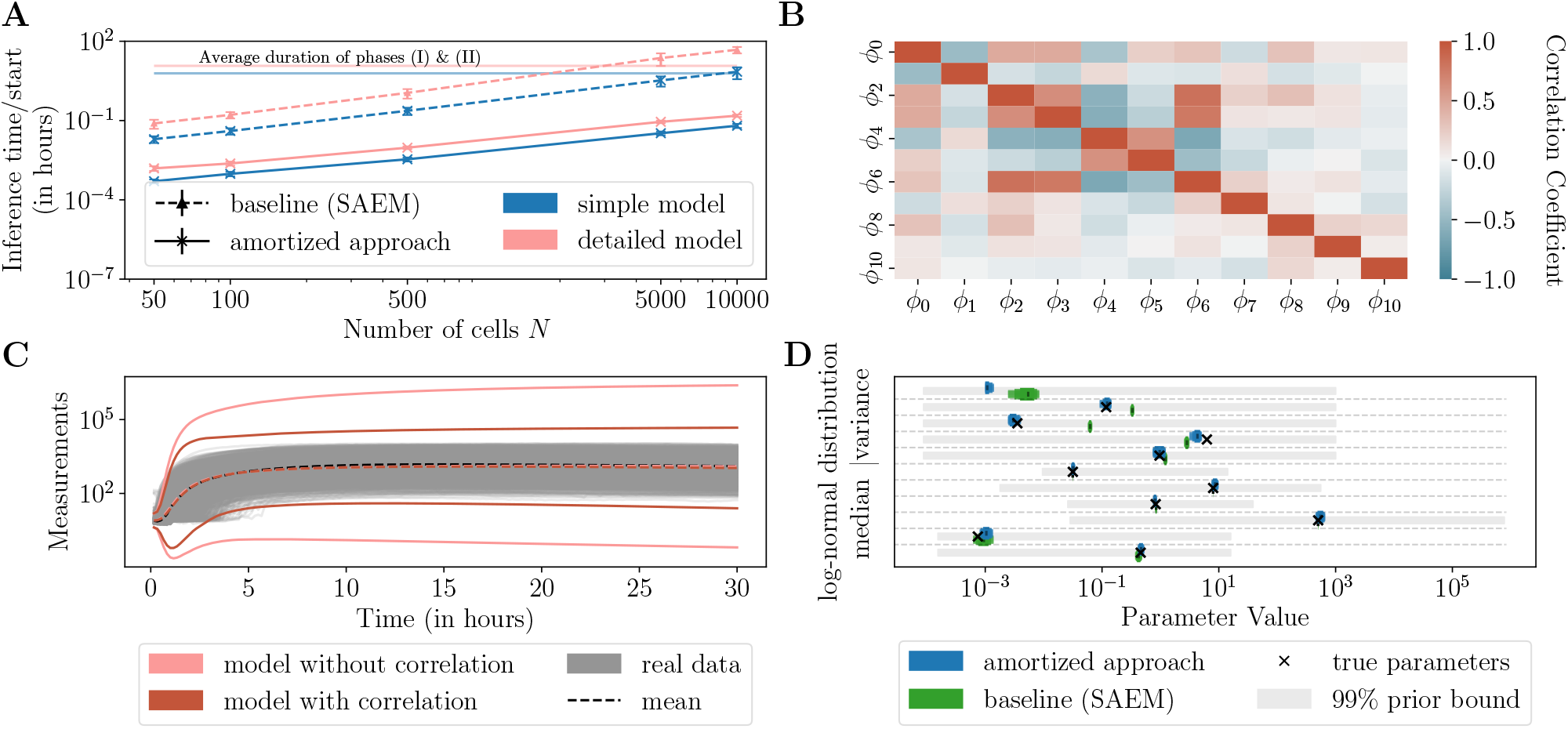
Flexibility and scalability of the amortized approach on the single-cell NLME models. (**A**) Computation time of the amortized inference phase (median *±* one standard deviation of a single run out of 100 multi-starts) for the single-cell NLME models compared to the baseline using parallelization. (**B**) Correlation of the posterior medians of the individual-specific parameters ***ϕ*** in the detailed single-cell model on real data. (**C**) Mean and 99% confidence intervals of the simulations for the detailed NLME model, where the population parameters are assumed to be log-normally distributed with and without correlations between parameters. (**D**) 80%, 90%, and 95% confidence intervals (CIs) for the simple single-cell NLME model (see Supplement Figure S15 for the other models) using synthetic data with known true parameters.

As described above, the parameters in the single-cell NLME models were assumed to be independently distributed. However, cross-correlations between parameters are essential to explain population behavior (Llamosi et al., 2016) but were not captured in (Fröhlich et al., 2018) due to computational costs. Indeed, for the detailed mRNA transfection model, the individual-specific posteriors of the respective parameters show a clear correlation (Figure 3B).

Therefore, we changed the population model to allow for a full covariance matrix of the random effects within each individual and repeated the amortized inference phase without further training of the neural posterior estimator. Including these correlations substantially improved the fit of the population variance (Figure 3C), which confirms the findings on the importance of incorporating cross-correlations between parameters in (Llamosi et al., 2016).

Beyond point estimates, we explored the possibility of performing accurate uncertainty quantification using a profile likelihood analysis, given the computational efficiency of the inference phase in our approach (Figure 3D and further details in Supplement A.4.4).

In summary, our analyses showed that our approach scales to large populations and allows for the reuse of the trained neural posterior estimator on different data sets and for different population models at almost no additional computational cost, rendering it substantially more scalable than SAEM.

### 3.3. Flexible generative model makes stochastic mixed-effects models easily tractable

As our approach based on neural posterior estimation proved to be valuable for deterministic models, we assessed its capability to cope with stochastic models, which can provide a more adequate description of the underlying process (Wilkinson, 2009; Stumpf et al., 2017). Ignoring the inherent stochastic nature of reactions at single-cell level can bias parameter estimates (Wiqvist et al., 2021), and pooling measurements from several cells is indispensable for reliable estimates (Zechner et al., 2014). Yet, the likelihood function is often unavailable for such stochastic models, which requires computationally demanding techniques such as approximate Bayesian computation or a Metropolis-within-Gibbs algorithm, which can handle the unavailable likelihood via correlated particle filters (Wiqvist et al., 2021; Botha et al., 2021; Sisson et al., 2018). However, our purely simulation-based approach does not need the likelihood function but only a generative model for simulation.

Here, we again considered the processes of mRNA transfection, but described by a stochastic differential equation (SDE) as proposed by (Pieschner et al., 2022) (detailed model specification in Supplement A.1). This model is superior for the description of individual cells and improves parameter identifiability (Pieschner et al., 2022), but has not been used so far in an NLME modeling framework.

The evaluation using the SDE NLME model on synthetic data revealed that our machine learning-based approach was indeed able to accurately recover the stochastic NLME model parameters (Supplement S10). Further analysis of synthetic data generated by the SDE NLME model showed that the simple ODE NLME model estimated parameters such that the variance of the population was three times greater than the true variance while for the stochastic NLME model, the variance is only 1.3 times greater (Supplement S17). Therefore, the stochastic model is capable of capturing the data more accurately. This underlines that a deterministic model can give erroneous results if it inadequately captures the underlying processes. The overall training time (7.5 hours) is comparable to the simple ODE model (6 hours), and the amortized inference phase remains highly efficient.

The simple ODE model of the mRNA transfection processes consists partly of a product of parameters *k· m*_0_*· scale*, where the individual parameters were not structurally identifiable, which means that not all parameters can be determined from the data (Fröhlich et al., 2018). However, the SDE model offers a more detailed representation encompassing the individual parameters *k, m*_0_ and *scale*. Only using our amortizing NLME framework, we were able to identify all parameters (Supplement S16).

In summary, our amortized approach enables the use of either a deterministic or a stochastic NLME model, whichever is more appropriate. This not only enables a more profound understanding of the actual mechanism, but can also improve model identifiability.

### 3.4. Amortization allows for full Bayesian analysis and more complex models

So far, we have considered inference problems with densely observed time points. Here, we turn to a model from pharmacokinetics introduced in (Diekstra et al., 2017; Yu et al., 2015) that describes the distribution of an angiogenesis inhibitor, a drug that inhibits the growth of new blood vessels, and its metabolite in a multi-compartmental model. Using data from 47 patients, including covariates such as weight, sex, and drug dosage regimes, we analyzed an ODE model with five states and 13 parameters. This model shows oscillatory behavior due to multiple dosing events (see Figure 4A). For comparison, we also explored FOCEI (Wang, 2008), which is arguably the most commonly used infer-ence method in pharmacokinetic modeling, implemented in `NONMEM` (Beal & Sheiner, 1980). Given FOCEI’s known tendency to converge to local optima, we performed 300 repeated parameter inference experiments (Pinheiro, 1994).

**Figure 4.**
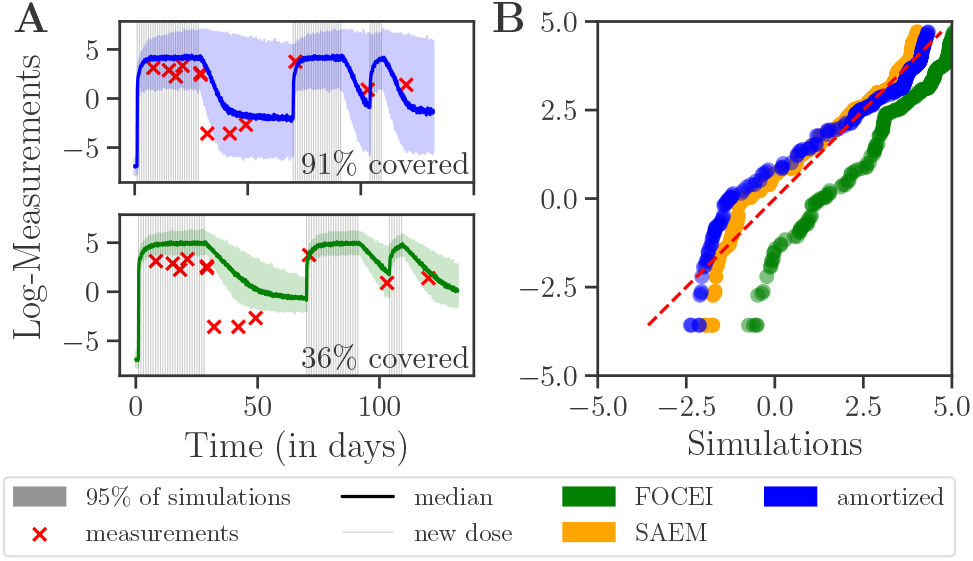
Baseline underestimates measurements. (**A**) Trajectory of sunitinib plasma measurements for one exemplary patient. Simulating samples from the population likelihood convoluted with the noise model using the covariates of this patient based on the estimated parameters of FOCEI and our amortized approach. (**B**) Ordered measurements (Sunitinib plasma) of all individuals against the median of 100 simulations per time point and individual.

Inferring population parameters yielded similar results for our amortized approach and SAEM. However, FOCEI underestimates the first observable consistently (see Figure 4B). Moreover, estimated variances with FOCEI are consistently smaller than those from our amortized approach or SAEM (see Supplement A.5). Simulating samples from the population likelihood convoluted with the noise model using estimated parameters from FOCEI and our approach, we found that only 72% (83%) of measurements fell within the region described by 95% (99%) of FOCEI’s estimated population variance. In contrast, our amortized approach covered 97% (99%) of measurements within the estimated variance (see also Figure 4A). Consequently, FOCEI systematically underestimates population variance – a known issue (Ge et al., 2004; Jönsson et al., 2004) – which is not encountered for the proposed approach.

Moreover, efficient evaluation of the population likelihood enables a full Bayesian analysis. By defining priors on population parameters, we can easily employ a Markov chain Monte Carlo method. Here, we used a Metropolis-Hastings sampler with adaptive proposal covariance as implemented in `pyPESTO` (Schälte et al., 2023). Generating 100, 000 samples takes only a few minutes, allowing us to analyze uncertainty in variance estimates. This analysis shows that FOCEI’s variance estimates do not fall within the 95% credible regions produced by the amortized approach, whereas the estimates of SAEM do. Yet, SAEM provides only a point estimate and approximate confidence intervals.

To show the applicability of our approach, we report the run times of the different methods. Running FOCEI repeatedly took 28 hours, while SAEM needed 37 hours. In contrast, our amortized approach, including repeating phase (III) 200 times and generating 100, 000 samples from the full population posterior, completed in 27 hours.

This demonstrates the feasibility of a full Bayesian analysis and highlights our method’s capability to handle complex models with individual-specific dosing regimes. Notably, we observed a less biased population fit compared to FOCEI.

## 4. Related Work

The inference methods for NLME models most commonly used today are deterministic, starting from the first inference method based on a first-order approximation of the model function around the expected value of random effects (Beal & Sheiner, 1980) and later on conditional modes (Lindstrom & Bates, 1990). The first-order approximation was used, among others, to analyze clinical patient data (Sheiner & Beal, 1980). Pinheiro & Bates reviewed more accurate methods based on the marginal likelihood approximation using Laplace methods or quadrature rules, obtaining potentially higher accuracy at higher computational cost (Pinheiro & Bates, 1995). Today, first-order conditional estimation with interaction (FOCEI) (Wang, 2008) is arguably the most common inference method used in pharmacokinetic modeling. Yet, the aforementioned methods have two main statistical drawbacks. First, they do not necessarily converge to the maximum likelihood estimates, and second, the estimates can be substantially biased when the variability of random effects is large (Ge et al., 2004; Jönsson et al., 2004).

For unbiased results, the stochastic expectation maximization algorithm (SAEM) was introduced, which converges under very general conditions (Kuhn & Lavielle, 2005). This method was applied, for example, to model the response of yeast cells to repeated hyperosmotic shocks (Llamosi et al., 2016). Yet, the algorithm can be computationally demanding, depending on the number of random effects and the structural complexity of the model.

However, all the methods mentioned do not apply to stochastic models for individuals, such as stochastic differential equations (SDEs). So far only Bayesian methods can provide exact inference and inherently facilitate uncertainty quantification for SDEs, but are even more computationally demanding (Wiqvist et al., 2021; Botha et al., 2021).

A simulation-based approach has been proposed to accelerate Bayesian inference in NLME models by approximating the likelihood of the population with simulated measurements and hand-crafted filters (Augustin et al., 2023). However, choosing an effective filter requires experimenting with different filters and numbers of simulations, while the efficiency depends on the availability of the gradient of the underlying simulator, which ours does not. Generalizing this to stochastic systems adds further complexity due to increased noise in the filter likelihood.

Further, one could directly learn the likelihoods based on neural likelihood estimation (Durkan et al., 2018). A different approach would be to directly tackle the full population likelihood or posterior. For example, Radev et al. (2020) used permutation embedding networks that can amortize over a given number of fixed i.i.d. trials, which could potentially be used in a setting with smaller data sets.

In general, computational costs make it difficult to fit NLME models to large data sets and to obtain reliable estimates of model parameters (Fröhlich et al., 2018; Augustin et al., 2023; Persson et al., 2022).

## 5. Discussion

We developed a novel approach for parameter inference in non-linear mixed-effects models based on amortized neural posterior estimation. The proposed method offers several advantages, such as scalability, flexibility, and accurate uncertainty quantification over established approaches, as we demonstrated on single-cell and pharmacology models.

One of the most important benefits of the method is its scalability. The efficient amortizing inference phase allows to scale to large numbers of individuals and can be applied to previously unseen data. The main computational bottleneck, the simulation and training phases, can be tackled by more extensive parallelization on a high-performance infrastructure since all simulations are independent. Further, the method can be applied to various population models and new data sets with low computational costs using the same trained neural posterior estimator, allowing efficient model selection. In contrast, state-of-the-art methods require a full restart for each population model.

Our machine learning-based approach is purely simulation-based; that is, it does not require the evaluation of likelihoods, but only a generative model to simulate synthetic data. Therefore, it can be used even for complex stochastic models, which established approaches fall short of, as we demonstrated on an SDE-based NLME model of mRNA transfection. Our approach can be easily extended to Markov jump processes, e.g., simulated with the Gillespie algorithm (Gillespie, 1977). This generality with respect to the generative model is unique in the NLME context, as special frameworks were needed to cope with stochastic differential equations (Wiqvist et al., 2021) or Markov jump processes (Zechner et al., 2014).

Lastly, the efficient neural posterior estimator facilitates more accurate and systematic methods to assess parameter uncertainty, here demonstrated by combining our approach with profile likelihoods and Bayesian inference. Conceptually, other uncertainty analysis methods, such as bootstrapping, could also be applied efficiently.

The study raises the question of how amortization can be best used in a hierarchical setup, in which, for example, the order in which problems need to be solved can be influenced. Furthermore, we consider it interesting to develop methods that assess on-the-flight whether problem-specific approximations, which are sequentially updated or an amortized approach is more beneficial for overall efficiency. This becomes relevant for small data sets and if no population model selection or accurate uncertainty quantification is needed since the computation time of the amortized approach will be higher compared to established methods.

Additionally, the proposed method may produce erroneous parameter estimates if the prior is too narrow or if the underlying model is misspecified (Schmitt et al., 2022), or use non-conservative posterior estimates (Hermans et al., 2022). However, misspecification of the model is a general problem for parametric methods. A solution might be to extend the loss function during training to include a misspecification measure (Schmitt et al., 2022). Furthermore, the accuracy of the approximated posteriors can be checked after training, e.g., by simulation-based calibration (Talts et al., 2020), or individual posterior checks by Markov chain Monte Carlo (MCMC) or approximate Bayesian computation (ABC) (Sisson et al., 2018). However, checking individual posteriors introduces an additional computationally expensive step.

Imperfect approximations of true posteriors can occur if the conditional normalizing flow, which might be the case for multimodal distributions (Hagemann et al., 2023). Then, the approximations could be improved by relaxing the constraints of the architecture imposed by invertible neural networks using generalized normalizing flows (Hagemann et al., 2023) or flow- and score-matching methods (Sharrock et al., 2022; Geffner et al., 2023; Dax et al., 2023). Nevertheless, we did not encounter such difficulties here, as we could approximate even the multimodal distributions in the simple ODE model. Thus, we are confident that our approach based on conditional normalizing flows can provide good estimates for the parameters in an NLME model.

In conclusion, the amortized approach we presented in this study offers a powerful solution for non-linear mixed-effects modeling. The approach enables researchers to flexibly use models for individuals and the population while performing accurate parameter estimation and uncertainty analysis in a more scalable manner than state-of-the-art methods.

## Code Availability

The code and a guide, aimed at assisting users in applying their own non-linear mixed-effects models, can be found at https://github.com/arrjon/Amortized-NLME-Models/tree/ICML2024. A snapshot of the code and the results underlying this study can be found at https://zenodo.org/record/8245786. The single-cell data has been made available by Fröhlich et al. (2018).

## Impact Statement

This paper presents a more efficient approach to parameter inference in non-linear mixed-effects models, which promises especially advances in medicine, drug development, and pharmacokinetics. While ethical considerations such as data privacy and equitable access should be prioritized for responsible implementation, increased efficiency contributes to resource conservation and highlights the potential for positive societal and environmental impacts.

## Acknowledgments

This work was supported by the German Federal Ministry of Education and Research (BMBF) (EMUNE/031L0293C), the German Research Foundation (DFG) under Germany’s Excellence Strategy (EXC 2047-390685813 and EXC 2151-390873048) and the project ID 458597554 - SEPAN, the Schlegel Professorship for J.H., and the EU Horizon 2020 program (ORCHESTRA; 101016167). Y.S. acknowledges financial support by the Joachim Herz Stiftung.

## A. Supplementary Information

**Figure S1.**
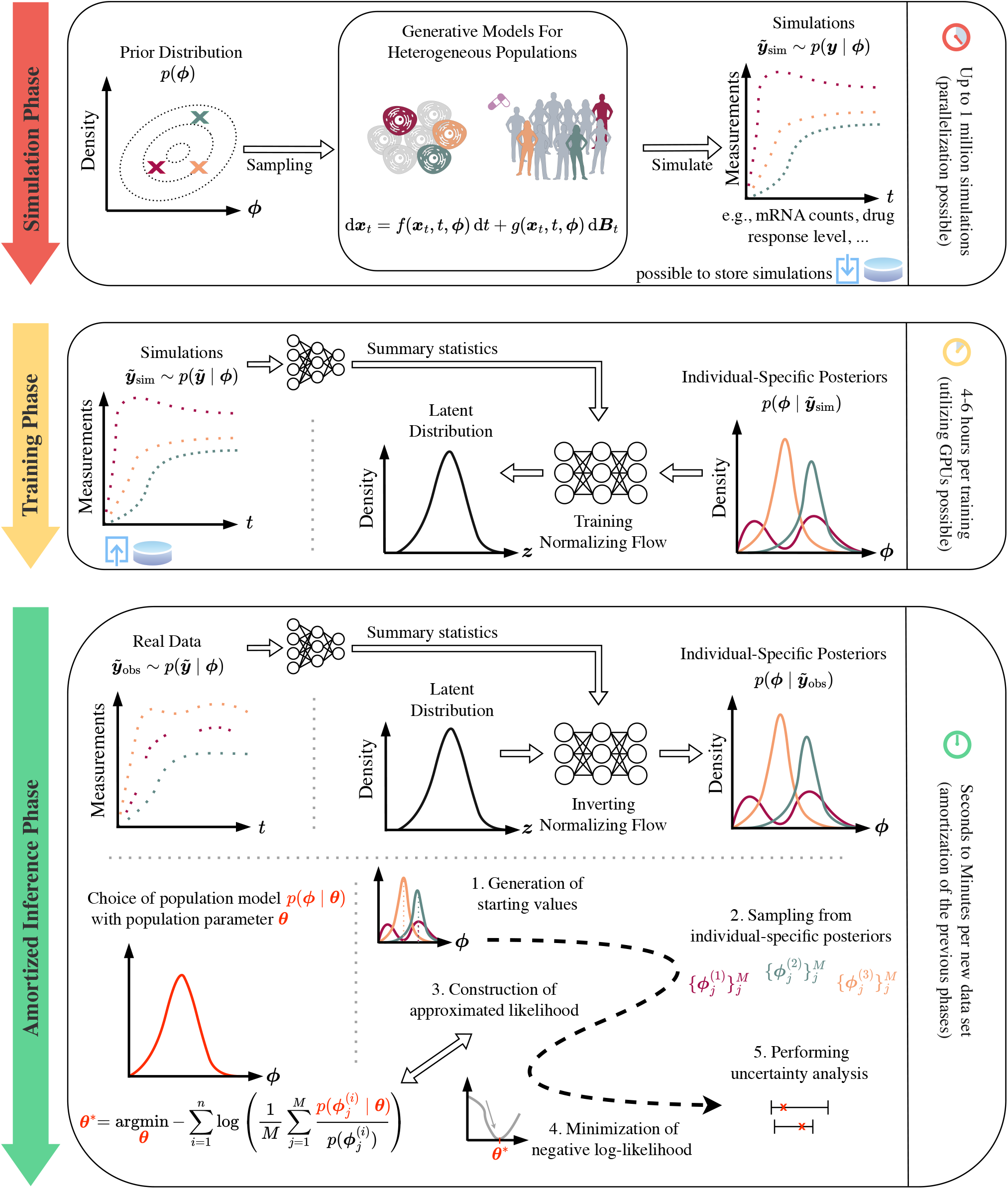
Detailed concept visualization of the neural posterior estimation based amortized approach to NLME model inference.

### A.1. Specification of the single-cell models

Living cells exhibit variability at the single cell level due to various factors such as mole processes, cell cycle state, environmental differences, and individual cell history (Fröhlich et al., 2018). Fröhlich et al. were interested in the dynamics of protein expression and transfected single cells with enhanced green fluorescent protein (eGFP)-encoding mRNA. The expression of the eGFP reporter gene was recorded every ten minutes for a period of 30 hours using a scanning time-lapse microscope setup. From these data, the authors estimated the parameters of the translation process using ordinary differential equation (ODE) models in an NLME framework.

In this work, we focus on two models termed the “simple” and “detailed” models from (Fröhlich et al., 2018). We denote the abundance of mRNA as *m*, proteins as *p*, ribosomes as *r*, and enzymes as *e*. For both models, we assume additive normal measurement noise, that is, the measurements 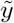 follow 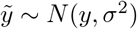 and our assumed prior distribution for *σ* is log *N* (−1, 2).

#### Simple ODE model

The ODE system has variables ***ϕ*** = (*δ, γ, km*_0_ *scale, t*_0_, *offset, σ*) and is defined as

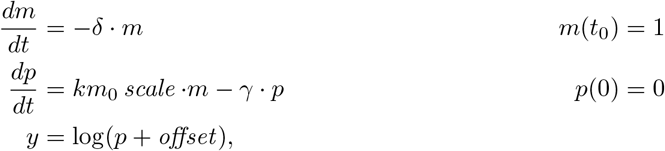

where the priors assumed for the variables are

- mRNA degradation rate *δ* ∼ log 𝒩 (−3, 5),
- protein degradation rate *γ* ∼ log 𝒩 (−3, 5),
- combined parameters *km*_0_ *scale* ∼ log 𝒩 (5, 11),
- mRNA entering the cell time point *t*_0_ ∼ log 𝒩 (0, 2),
- and *offset* ∼ log 𝒩 (0, 6).

The parameters *k, m*_0_, *scale* can only be identified as a product to improve identifiability (Fröhlich et al., 2018). This ODE system has an analytical solution, which we use to perform simulations in Python.

#### Detailed ODE model

The ODE system has variables ***ϕ*** = (*δ*_1_*m*_0_, *δ*_2_, *e*_0_*m*_0_, *k*_2_*m*_0_ *scale, k*_2_, *k*_1_*m*_0_, *r*_0_*m*_0_, *γ, t*_0_, *offset, σ*) and is defined as

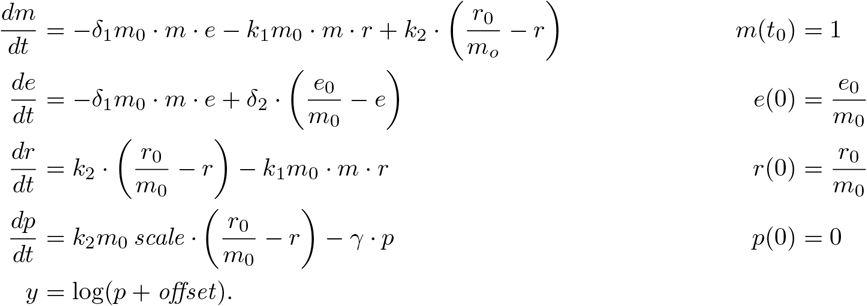

For a detailed description of the parameters, we refer to (Fröhlich et al., 2018). The priors assumed for the variables are

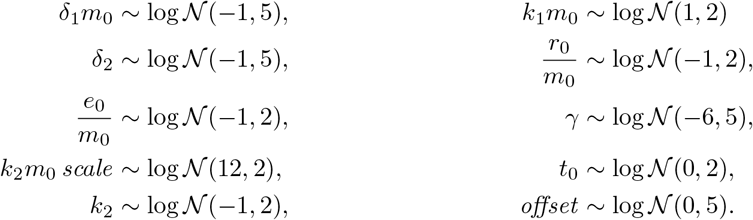

This ODE system is simulated using the Rodas5P solver implemented in the Julia package `DifferentialEquations.jl` (Rackauckas & Nie, 2017).

#### SDE model

The simple ODE model can be easily extended to the SDE model

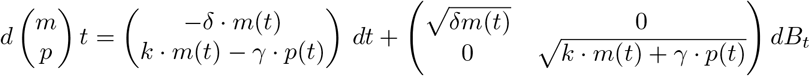

from (Pieschner et al., 2022) with ***ϕ*** = (*δ, γ, k, m*_0_, *scale, offset, σ*), where *B*_*t*_ is a two-dimensional standard Brownian motion, *m*(*t*_0_) = 1 and *p*(0) = 0. To compare the model to the previous one we take as observable mapping

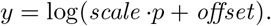

The priors assumed for the variables are

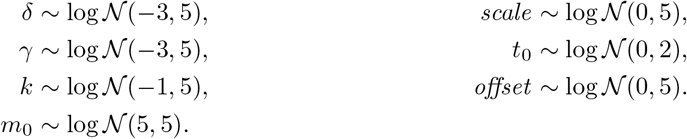

This SDE system is simulated based on an Euler-Maruyama scheme with a step size of 0.01 and using just in time compilation from `numba` (Lam et al., 2015).

##### A.1.1. Synthetic data

The synthetic data set is generated by setting the population parameters to reasonable values based on the results in (Fröhlich et al., 2018) (see Table 1, 2 and 3) and then sampling random effects from a log-normal distribution until the desired number of synthetic cells is generated. Since we know all cell-specific parameters, we can compute the sample mean and covariance of the parameters, which are the optimal values that we would like to recover. Fixed effects are modeled as random effects with 0 variance.

**Table 1.** Population parameters of log-normal distribution for synthetic data of simple single-cell NLME model.

| parameter | $\delta$ | $\gamma$ | $k m_0 \text{ scale}$ | $t_0$ | $\text{offset}$ | $\sigma$ |
| --- | --- | --- | --- | --- | --- | --- |
| mean | -0.694 | -7.014 | 6.217 | -0.164 | 2.079 | -3.454 |
| variance | 0.941 | 7.014 | 0.004 | 0.116 | 0 | 0 |

**Table 2.**
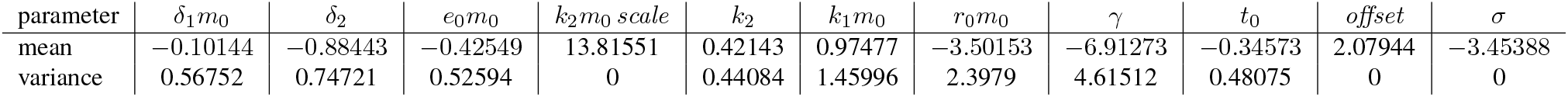
Population parameters of log-normal distribution for synthetic data of detailed single-cell NLME model.

**Table 3.** Population parameters of log-normal distribution for synthetic data of the SDE single-cell NLME model.

| parameter | $\delta$ | $\gamma$ | $k$ | $m_0$ | $scale$ | $t_0$ | $offset$ | $\sigma$ |
| --- | --- | --- | --- | --- | --- | --- | --- | --- |
| mean | -0.694 | -7.014 | 0.027 | 5.704 | 0.751 | -0.164 | 2.079 | -3.454 |
| variance | 0.941 | 7.014 | 0.675 | $6 \cdot 10^{-5}$ | 0 | 0.116 | 0 | 0 |

**Table 4.** Detailed runtime analysis. We report duration of the simulation phase, the average training time, and the number of simulations of the underlying ODE or SDE model. For SAEM we report the average number of ODE model simulations (assuming one simulation per individual per iteration) for the data set with 10, 000 cells and for all data sets and repeated inference runs combined. For any new data set, there are no additional simulations in the amortized approach. SAEM is not capable to perform inference for the SDE NLME model.

| model | (I)<br>simulation phase | (II)<br>training phase | epochs | total<br>simulations | # simulations in<br>one SAEM run | total number of<br>simulations in SAEM |
| --- | --- | --- | --- | --- | --- | --- |
| simple | - | 6.11 h | 416 | 53 Mio. | 34.8 Mio. | 5 030 Mio. |
| detailed | 3.74 h | 8.00 h | 355 | 64 Mio. | 99.8 Mio. | 9 640 Mio. |
| SDE | 2.16 h | 5.12 h | 417 | 64 Mio. | - | - |

### A.2. Implementation

We implemented the individual-specific posterior approximation using the `BayesFlow` tool (Radev et al., 2023). To estimate the population parameters, we implemented the optimization problem (5) as an objective function in the `pyPESTO` toolbox (Schälte et al., 2023). There, we used the local optimization method L-BFGS (Liu & Nocedal, 1989) implemented in `SciPy` (Virtanen et al., 2020) embedded in a multi-start framework with uniformly sampled starting points in the 99% range of the prior. In our applications, usually ten starts were already enough to reliably obtain the same optimum several times (see Figure S14). Parameters that are shared between individuals, that is, parameters that do not consist of a random effect, can be approximated in the given approach by fixing their variance to a small value.

As invertible neural networks, we used neural spline layers (Durkan et al., 2019) with variable depth of seven to eight layers and two to three coupling layers. Since all models describe trajectories over time, we chose a long short-term memory (LSTM) network (Hochreiter & Schmidhuber, 1997) with 2^*d*^ units as our summary network. We choose *d* such that the number of units is larger than the number of observations given by the model.

For each model, multiple neural posterior estimators were trained. We varied the number of invertible layers from six to eight, added a 1*d*-convolutional layer on top of the LSTMs and a dense layer at the end. Training consists of several epochs, and in each one we generated 1000 batches of 128 simulations. Simulations can be generated before or during training. Depending on the simulation time of the model, pre-simulation or online training is more efficient. We used online training for the simple ODE model, while we generated simulations beforehand for the other models. For all models, we set a maximum of 500 epochs and training was stopped earlier if the loss calculated on a validation set did not improve for five epochs, which reduced training time and is assumed to improve the generalization capacity of neural networks (Zhang et al., 2021). The error calculated on a validation set during training suggested convergence for all models (Figure S2).

For each start in `Monolix` (Lixoft SAS, 2023), we increased the iteration limit for each task (SAEM, standard error, and likelihood estimation) to ensure convergence at each step. We set the maximum number of iterations in the two phases (exploratory and smoothing) of SAEM to 10,000 and 1,000, respectively. `Monolix`’ auto-stop criteria usually stopped the algorithm before it could perform that many iterations. The iteration limit for estimation of the standard errors was set to 1,000 and the Monte Carlo size for likelihood estimation was set to 50,000. All other settings were left at their default values. The starting points were sampled from the prior.

For comparison, we also explored a multi-start approach using FOCEI (Wang, 2008), implemented in `NONMEM` (Beal & Sheiner, 1980), which is arguably the most widely used inference method in pharmacokinetic modeling. We performed 300 starts, where we sampled the starting values from the prior.

We ran all analyses on a computing cluster using eight CPU cores for parallelization and one GPU for training the neural networks. The computing cluster uses an AMD EPYC 7F72 CPU with a core clock speed of up to 3.2 GHz and 20 GB of RAM per available core. The neural network training was performed on a cluster node with an NVIDIA A100 graphics card with 40 GB of VRAM.

The simulations of the generative model, multi-starts in `pyPESTO` and a single start in `Monolix` used all available cores for parallelization. Moreover, the contribution of each individual could also be evaluated in parallel, giving the option of further parallelizing the calculation of the objective in a single start.

### A.3. Conditional normalizing flows provide accurate and efficient approximation of individual-specific posteriors

We checked the convergence of neural posterior estimators based on their calibration plots, a diagnostic tool that comes with `BayesFlow`. Simulation-based calibration is a method to detect systematic biases in any Bayesian posterior sampling method (Talts et al., 2020). Incorrect calibration can be seen by deviations from uniformity. None of our estimators showed systematic biases (Figure S3). Furthermore, for the best estimators, we assessed the validity of the individual-specific posteriors of the real data by comparing them with the posterior approximations given by an MCMC approximation with adaptive parallel tempering implemented in `pyPESTO` (Schälte et al., 2023). In particular, the bimodal distributions of the parameters *δ* and *γ* in the simple ODE model are nicely recovered (Figure S4). For the detailed model, MCMC showed poor convergence behavior over repeated runs with a small effective sample size.

**Figure S2.**
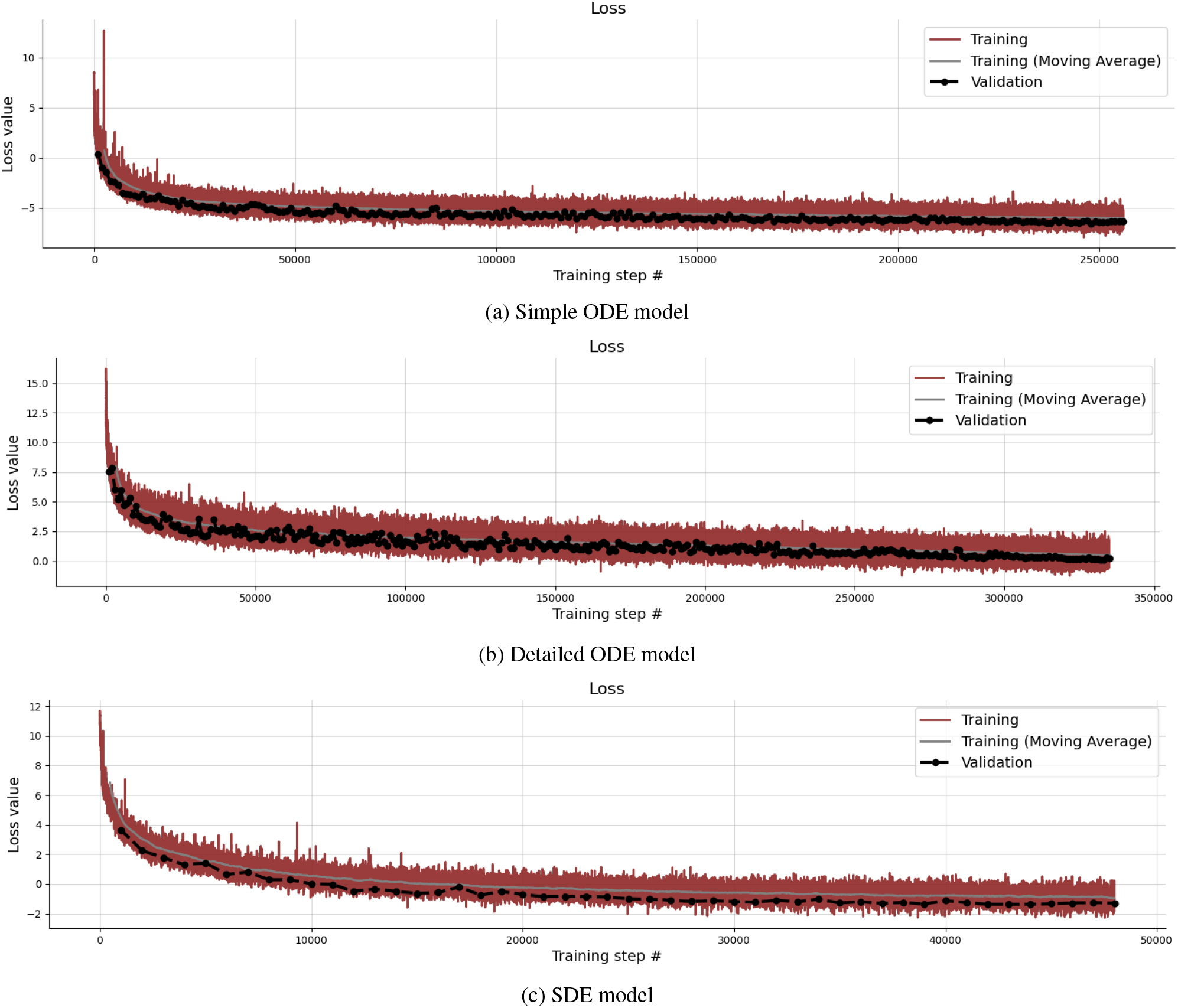
Exemplary loss during training.

An assessment of computation time revealed that the employed MCMC sampler required approximately 1 million samples and 10 chains with an effective sample size of 195, which took around 20 minutes of computation time for a single cell. In comparison, the trained neural posterior estimator only required a few seconds for the same effective sample size and on the same set-up (see details on the implementation in Methods A.2). Thus, in this case, the training time of the neural networks to obtain individual-specific posteriors, ∼ 6.5 hours, would be amortized after around 20 cells, or even after an individual cell if a sufficiently high sample size is required. This demonstrates the efficiency of neural posterior estimation for parameter estimation also outside a mixed-effects context.

**Figure S3.**
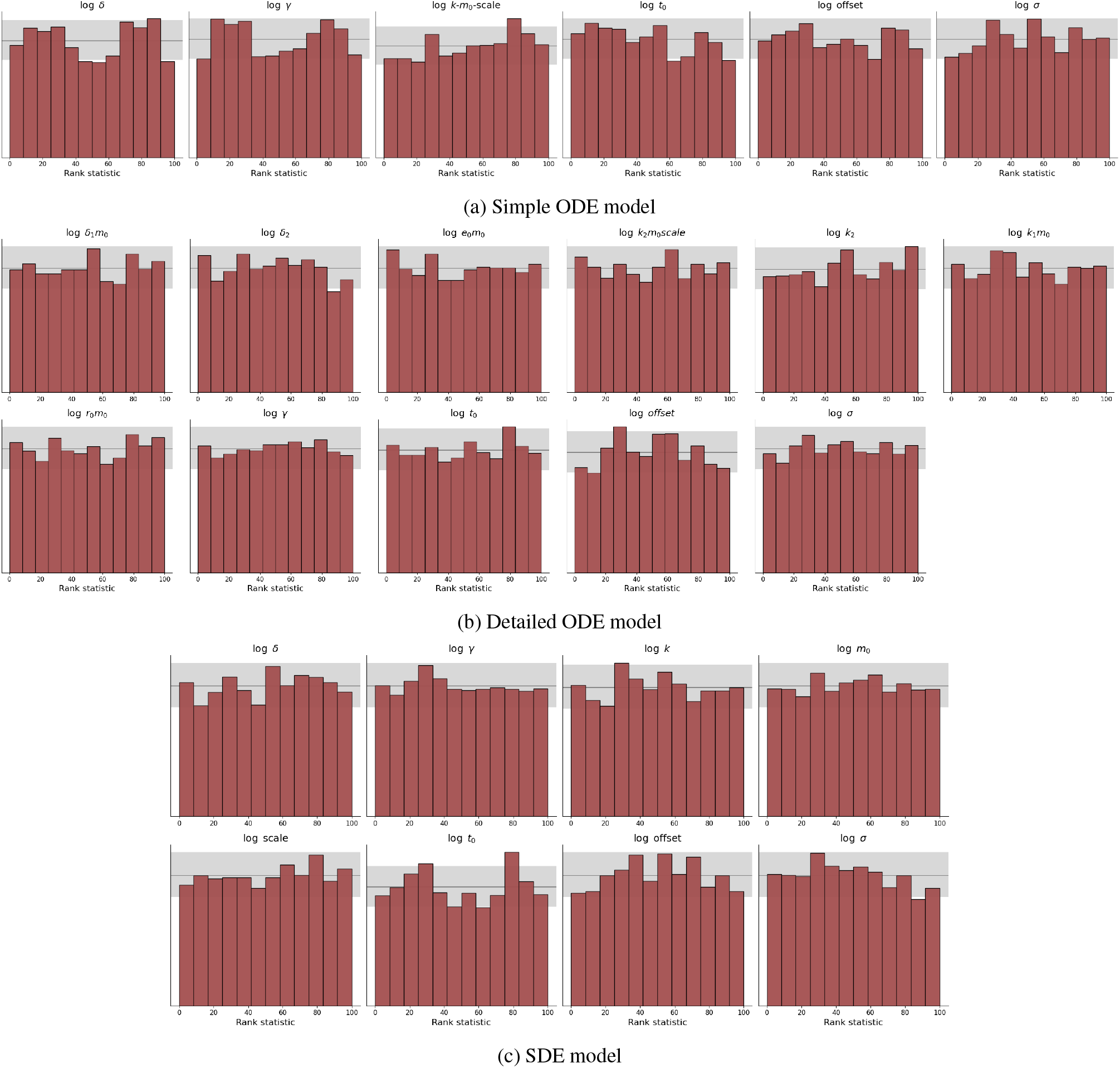
Simulation-based calibration plots of the individual posteriors for the (a) simple ODE, (b) detailed ODE and (c) SDE models. Incorrect calibration can be seen by deviations from uniformity (bars outside the gray area).

**Figure S4.**
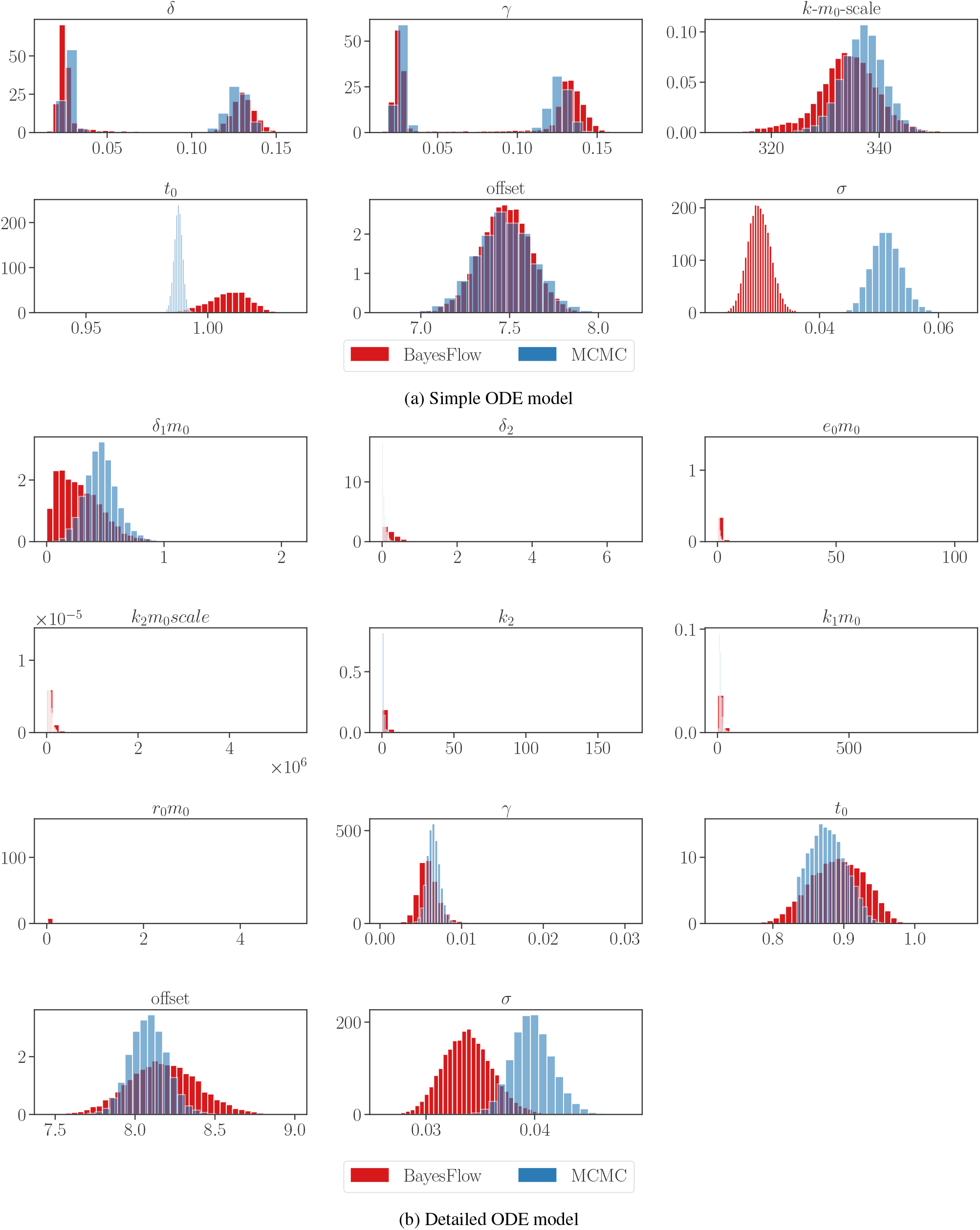
Comparing individual-specific posteriors from an MCMC approximation and the neural posterior estimator for a single real cell in the (a) simple and the (b) detailed ODE model.

**Figure S5.**
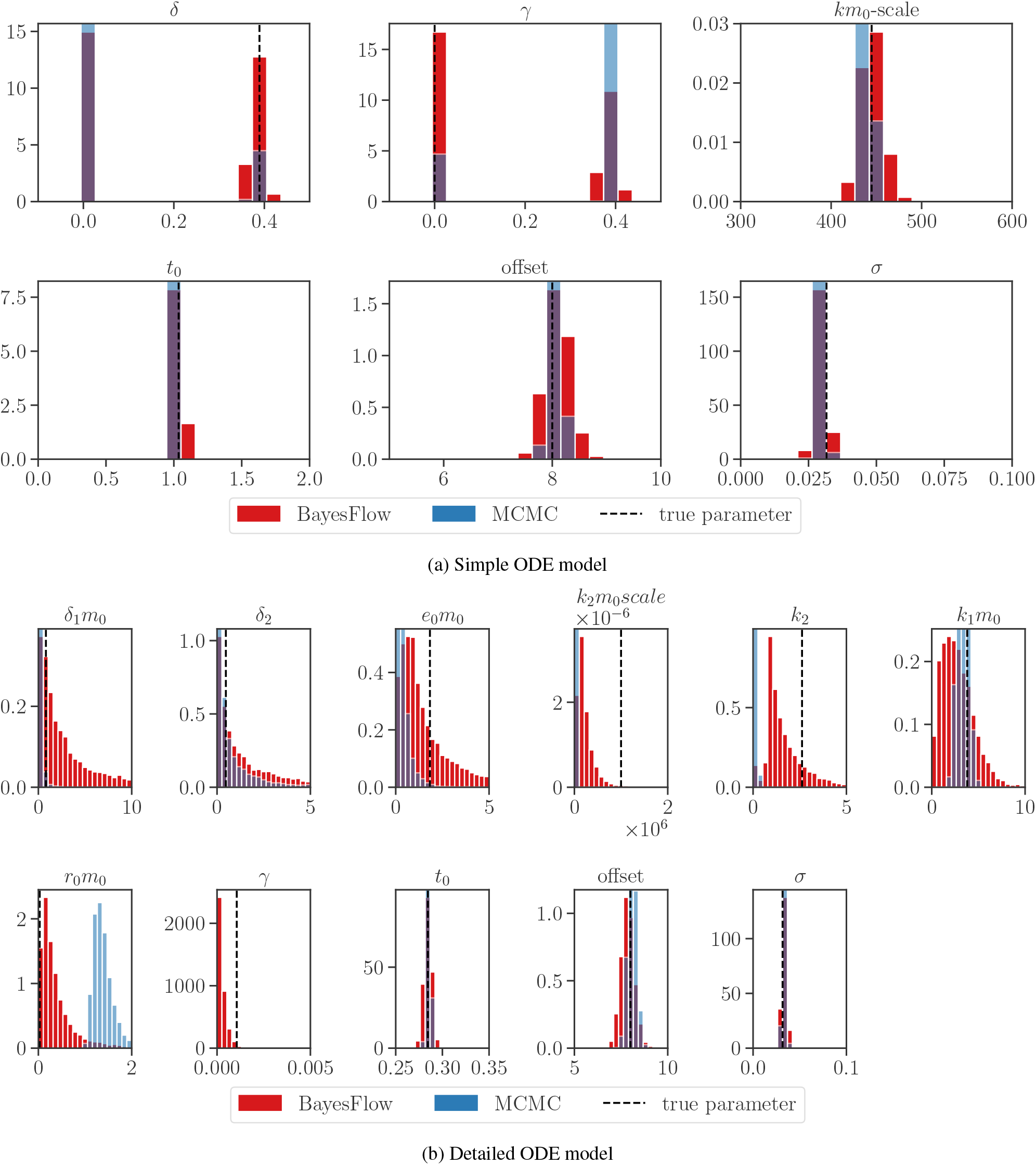
Comparing individual-specific posteriors from an MCMC approximation and the neural posterior estimator for a single synthetic cell in the (a) simple and the (b) detailed ODE model.

**Figure S6.**
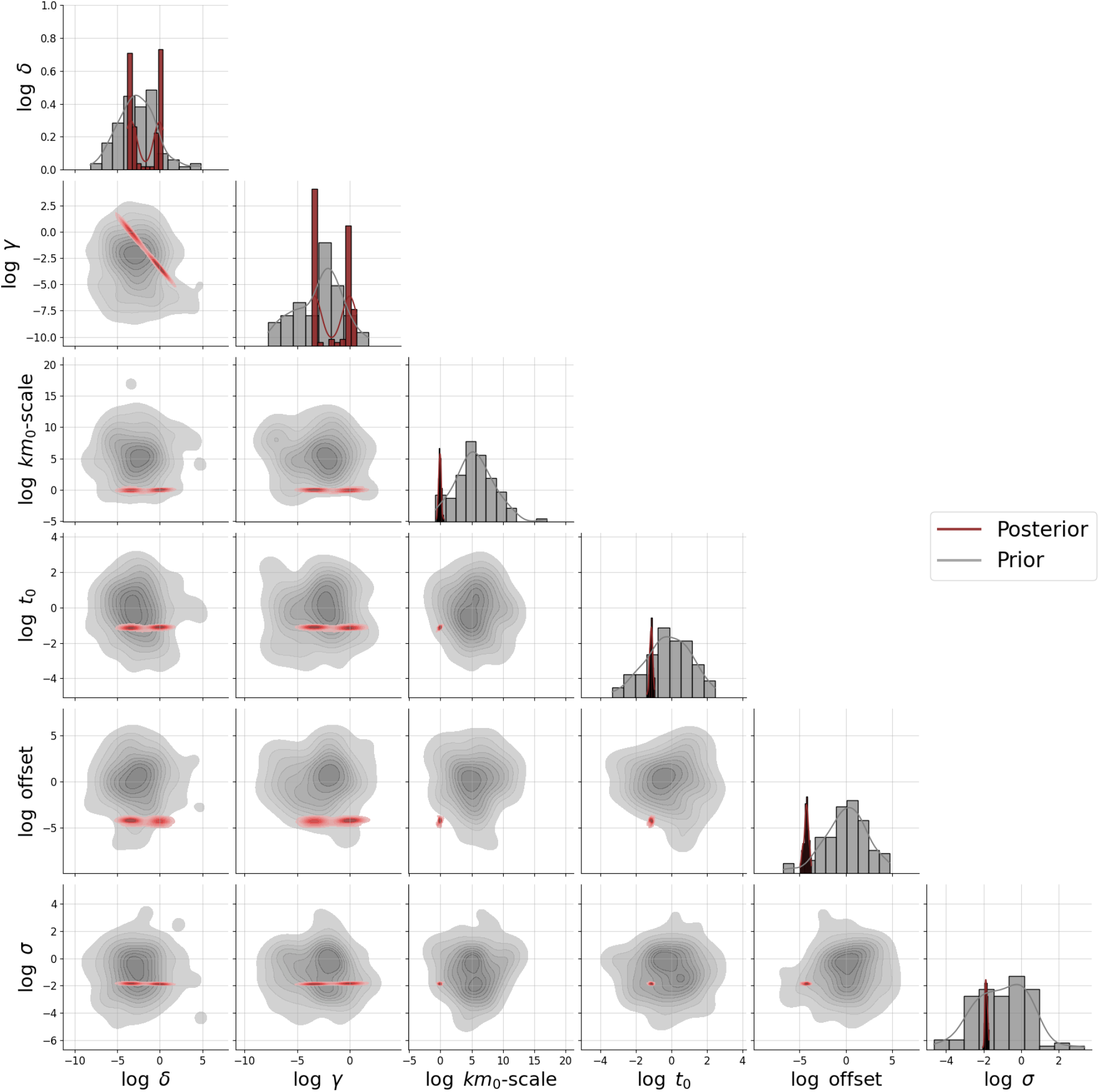
Full posterior for the simple ODE model and an exemplary real single cell

**Figure S7.**
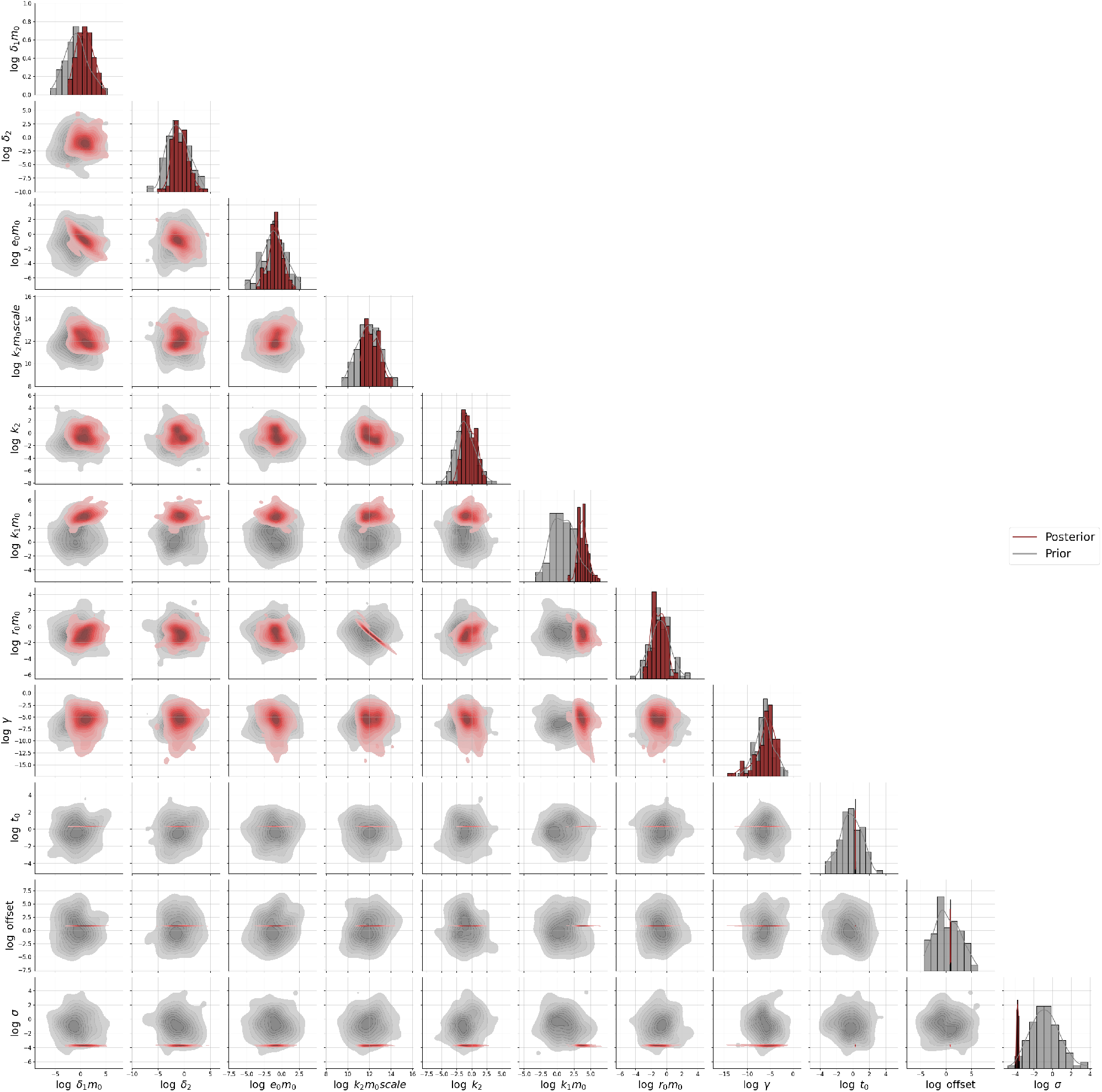
Full posterior for the detailed ODE model and an exemplary real single cell

**Figure S8.**
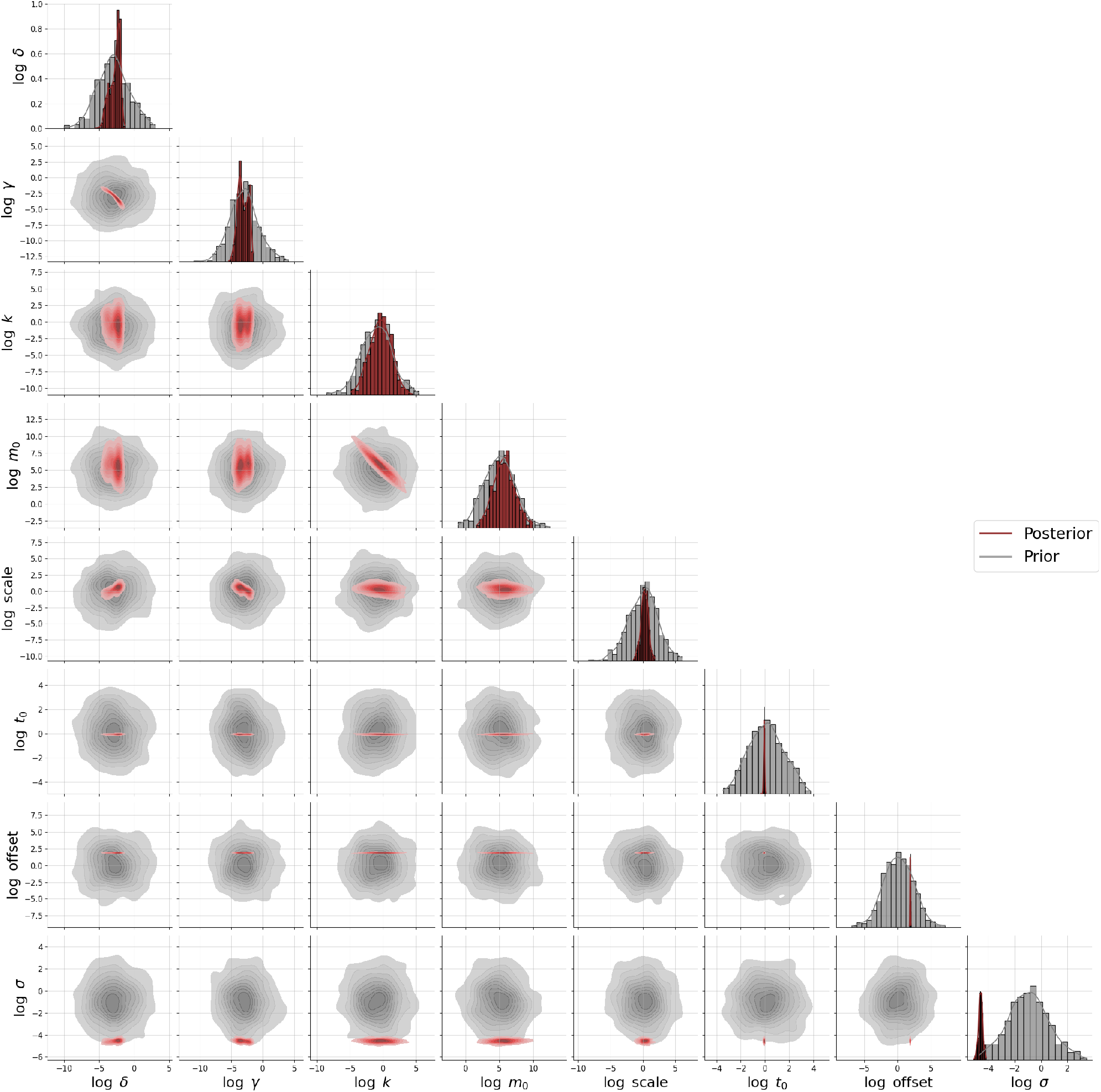
Full posterior for the SDE model and an exemplary real single cell

### A.4. Further analysis of the single-cell NLME models

**Figure S9.**
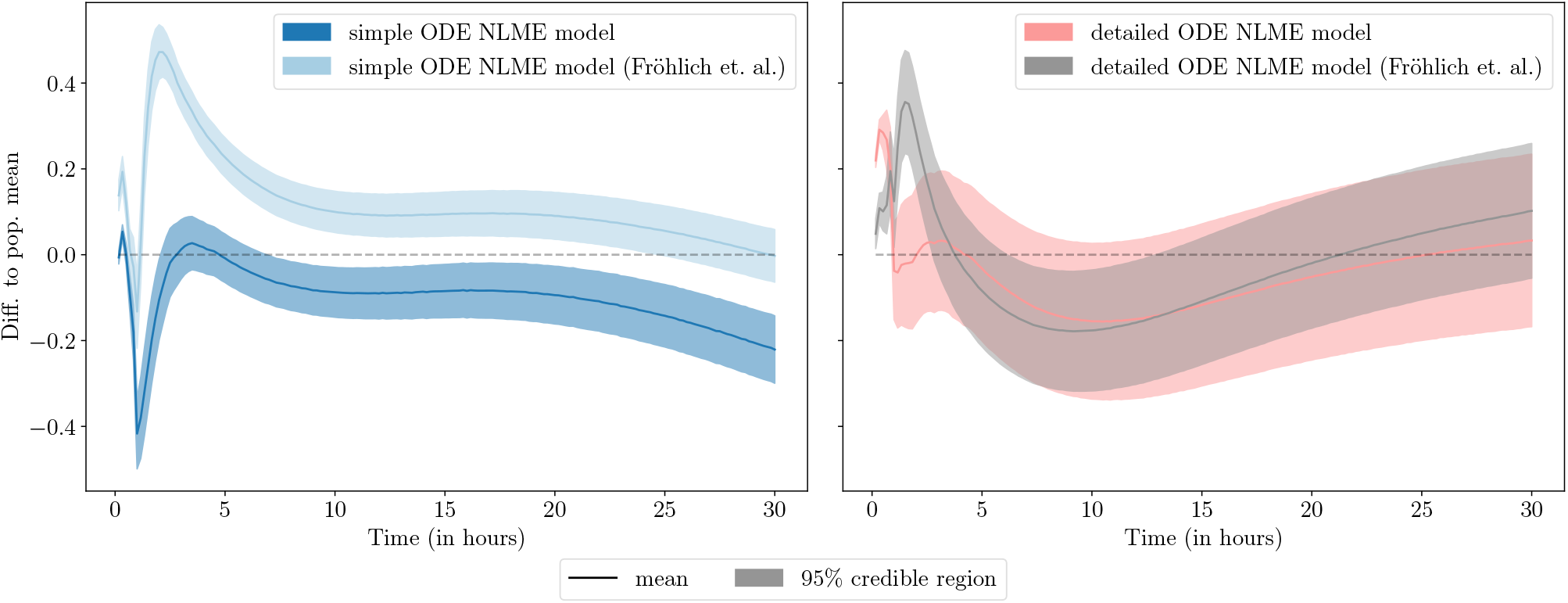
The difference in the population mean estimated from real trajectories and simulations generated with the estimated population parameters is shown with a 95% confidence interval (CI). The detailed model captures the population mean after 0.82 hours and then shows no significant deviation from the population mean, while the simple model needs 1.97 hours to recover the population mean and then differs significantly from the population mean. In addition to the models fitted with the amortized approach, the best fit of Fröhlich et al. (2018) for the simple and detailed ODE NLME model is shown (Fröhlich et al., 2018).

#### A.4.1. Influence of the number of posterior samples

We also checked the impact of the number of posterior samples *M* on the estimated population parameters. In Figure S10a, we see a small decrease in the mean squared error for a larger number of posterior samples across all ODE NLME models and data sets. Monte Carlo integration theory suggests that the error rate 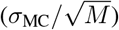 should be 5 times lower when increasing the sample size from *M* = 10 samples to *M* = 250. However, the median error decreases only by a factor of around 1.5. We assume that the lower rate comes from the fact that we already have a good approximation with a small number of samples due to the following observation: In the cases shown here, the population density *p*_pop_ and the prior *p*(***ϕ***) come from the exponential family. Hence, the loss function consists of logarithms of sums of exponentials. For example, if both are normal distributions 𝒩 (***β*, Ψ**), 𝒩 (***μ*, Σ**) respectively, then we can define 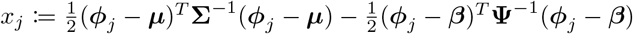. We can bound the logarithm of sums of exponentials, by the maximum function (Boyd & Vandenberghe, 2004) through

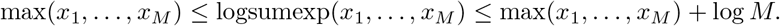

This is true since 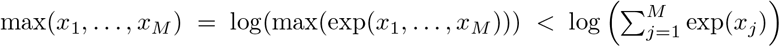 and 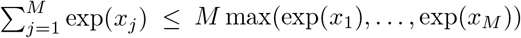. The latter inequality is an equality if and only if all *x*_*j*_ are equal. The log-sum-exp function is even convex (Boyd & Vandenberghe, 2004) and can be numerically stable evaluated by using the log-sum-exp-trick (Blanchard et al., 2021).

Therefore, the main contribution to the loss function comes from “good” posterior samples ***ϕ***_*j*_ that optimally balance the population distribution and individual priors (a maximal *x*_*j*_). If we add a “bad” sample to a set of already “good” samples, only the upper bound will change by log((*M* + 1)*/M*). Intuitively, this explains why we only need a small number of “good” samples to get a reasonable approximation of the population likelihood.

In Figure S10b, we see that the inference time increases when a larger sample size is used for the posterior, but that inference is still faster than the baseline method. Moreover, for each model, we reused the same neural posterior estimator on all data sets, whereas for the baseline method, we needed to restart the whole optimization. Therefore, to get estimates for 100 runs on multiple data sets, we saved 128 times of computational resources using the amortized approach compared to SAEM.

In Figure S11–S13, we see that for the amortized approach the variance of the parameter estimates of a multi-start optimization decreases with increasing posterior sample size or size of the data set. However, in general, for increasing size of the data set, more multi-starts were needed for our approach and SAEM (Figure S14). The amortized approach shows that for a posterior sample size of at least *M* = 50, most runs reach a similar likelihood value, in particular for the smaller data sets. Furthermore, the amortized approach is able to recover the multi-modality in the first two parameters of the simple ODE NLME model (Figure S11). This modality comes from the fact, that we can swap *δ* and *γ* in the simple ODE model without changing the solution of the ODE.

**Figure S10.**
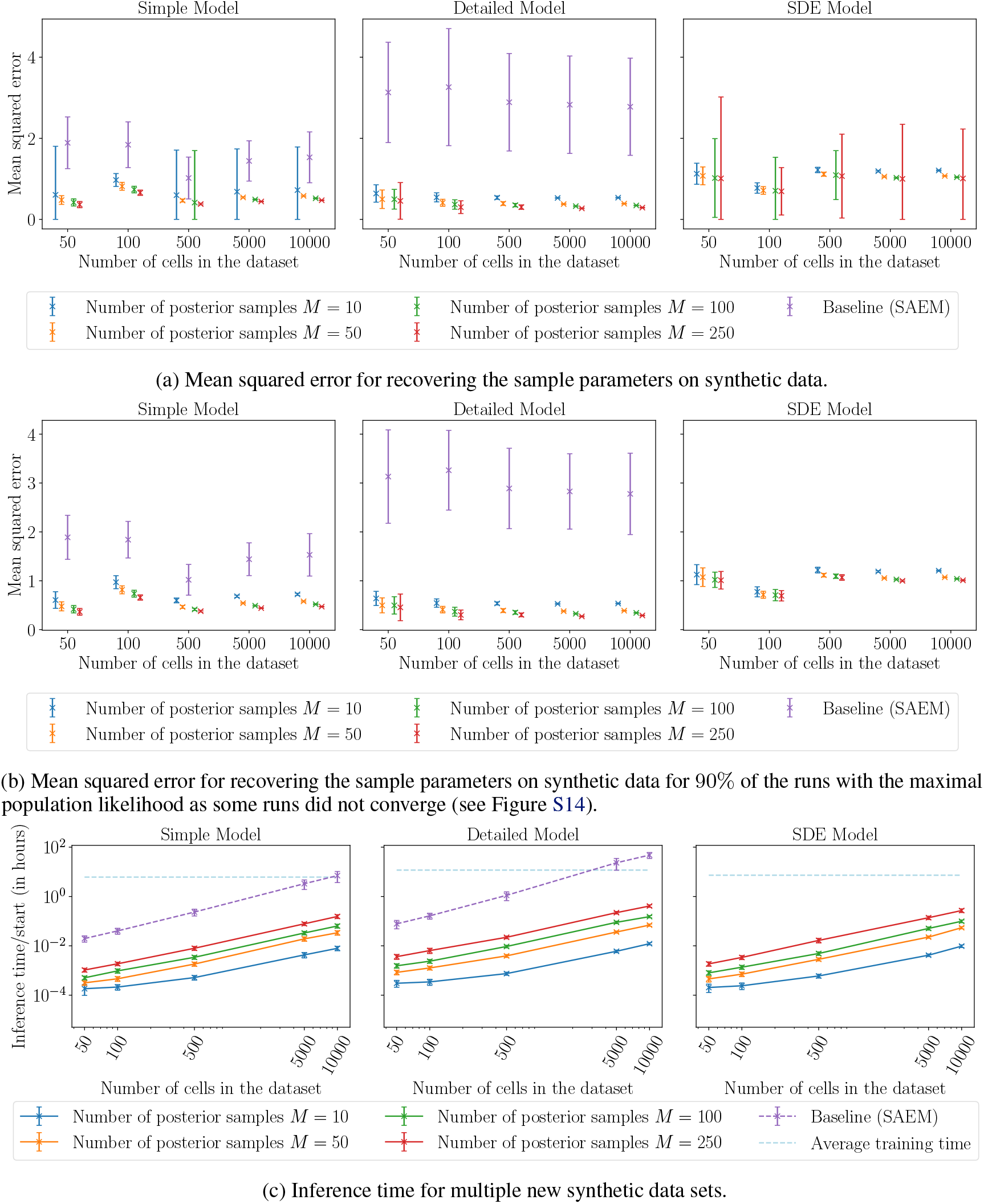
Accuracy and inference time for multiple new synthetic data sets for the single-cell NLME models. The median of the mean squared error for 100 runs is shown with one standard deviation. Different numbers of posterior samples were used to estimate population parameters. For each model, we reused the same neural posterior estimator on all data sets.

**Figure S11.**
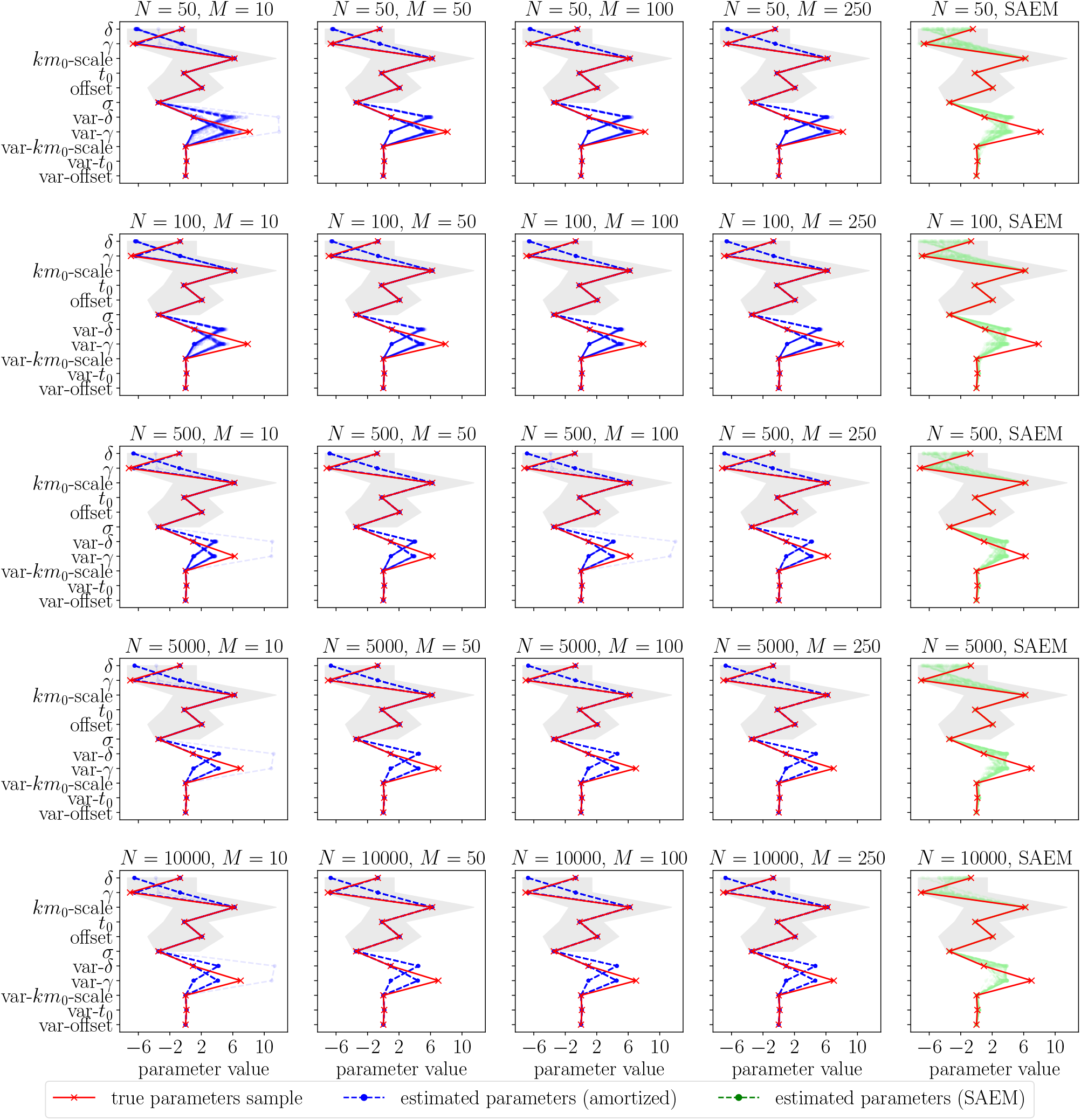
Parameter estimates on synthetic data sets for the simple single-cell NLME model for 100 runs. Different numbers of posterior samples *M* were used to estimate population parameters. For each data set of size *N*, we reused the same neural posterior estimator.

**Figure S12.**
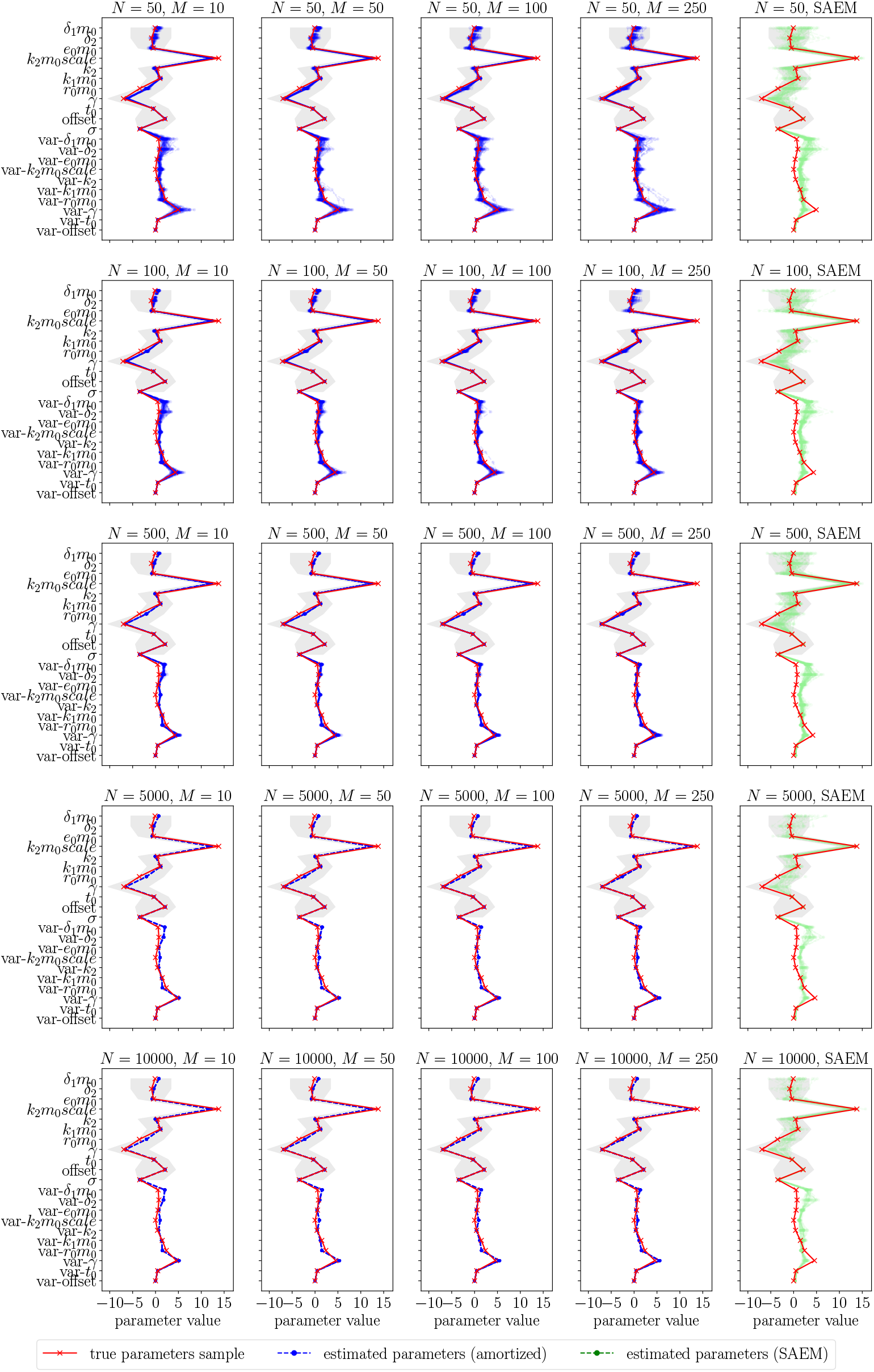
Parameter estimates on synthetic data sets for the detailed single-cell NLME model for 100 runs. Different numbers of posterior samples *M* were used to estimate population parameters. For each data set of size *N*, we reused the same neural posterior estimator.

**Figure S13.**
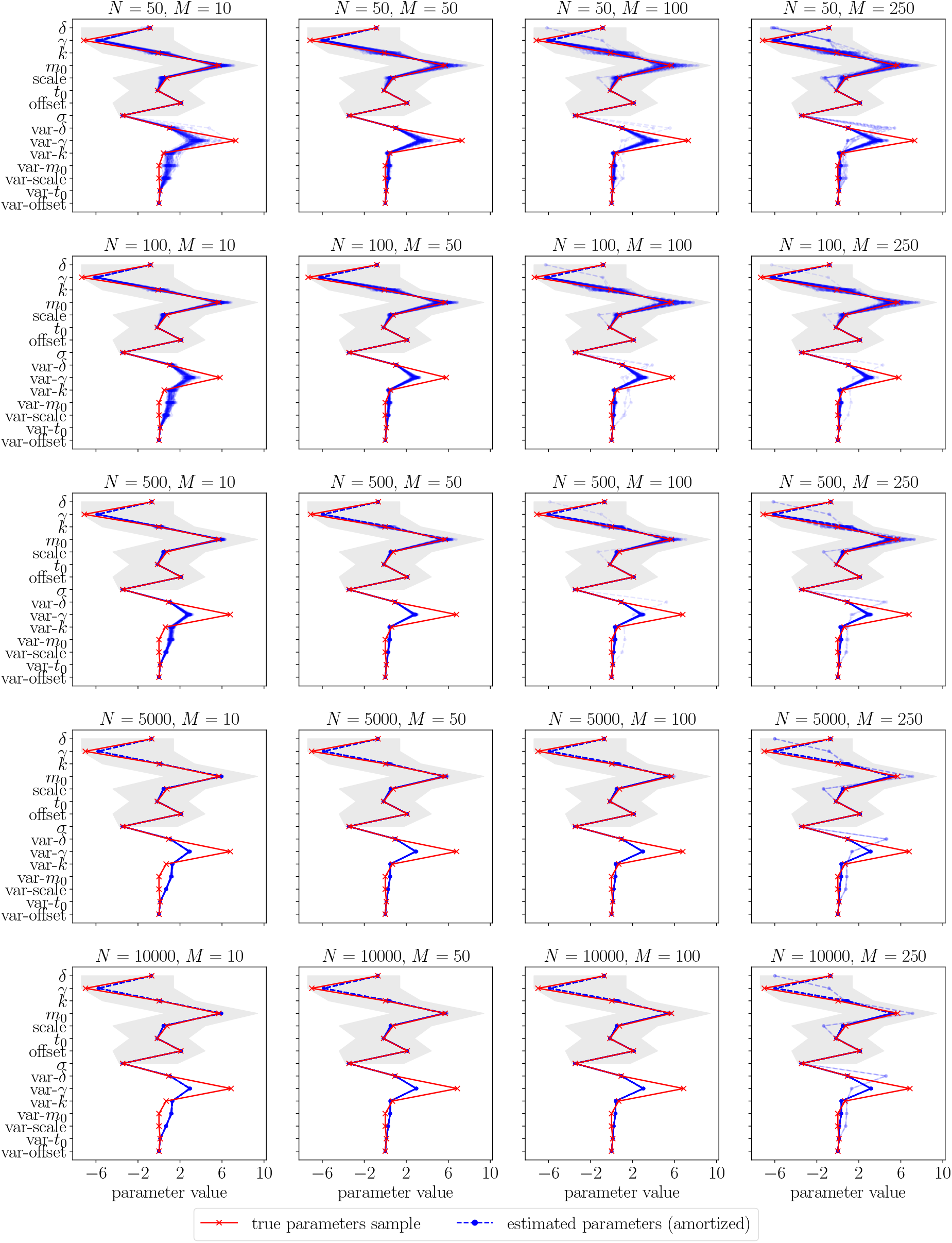
Parameter estimates on synthetic data sets for the SDE single-cell NLME model for 100 runs. Different numbers of posterior samples *M* were used to estimate population parameters. For each data set of size *N*, we reused the same neural posterior estimator.

**Figure S14.**
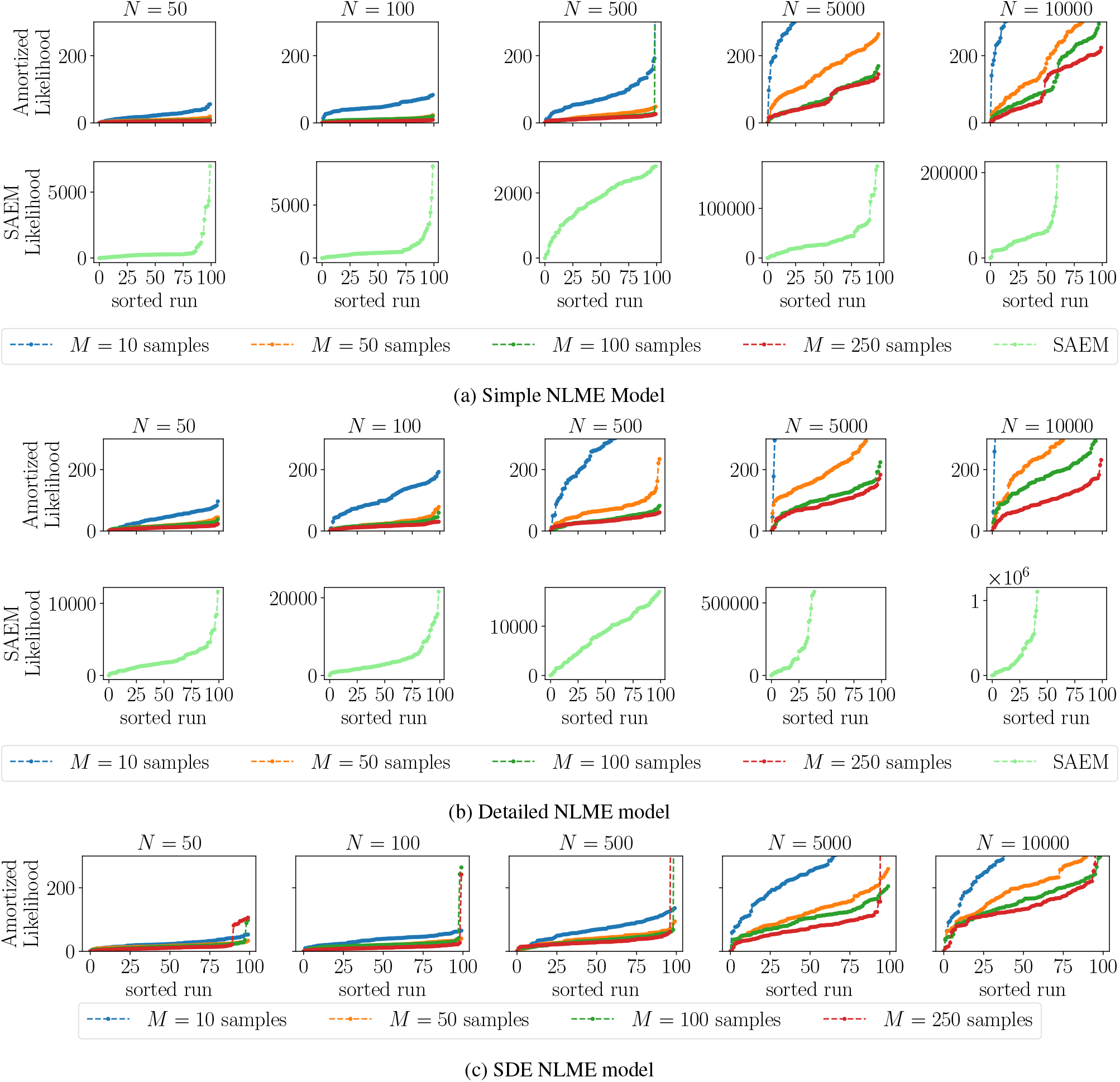
Approximated negative log-likelihood values (with an offset of the minimum value) on synthetic data sets for the single-cell NLME models for 100 runs. Different numbers of posterior samples *M* were used to estimate population parameters. For each data set of size *N*, we reused the same neural posterior estimator.

#### A.4.2. Uncertainty Quantification

#### A.4.3. Efficient inference of the population parameters enables robust uncertainty analysis

Our approach based on amortized neural posterior estimation allows efficient construction of point estimates. Beyond point estimates, in many applications, it is important to assess the uncertainty of the parameters to determine the identifiability of the parameters, draw reliable conclusions, and perform representative predictions (Raue et al., 2013; Maier et al., 2020). The implementation of SAEM in `Monolix` allows standard errors to be obtained through linearization of the likelihood or by a stochastic approximation of the Fisher information matrix, which yields asymptotically correct results under the assumption of normally distributed errors and a large amount of data. Using these standard errors, the confidence intervals are calculated using the Wald statistic (Lixoft SAS, 2023).

However, to ensure the validity of the confidence intervals, it is often advisable to use bootstrapping or non-local approaches such as profile likelihoods, as these are more accurate when the above assumptions are not met. This can, for example, allow for non-symmetric confidence intervals (Fröhlich et al., 2014). Such tests are infeasible when the computational time is high, as it is often the case with SAEM.

We explored the possibility of performing accurate uncertainty quantification, given the computational efficiency of the inference phase in our approach. Specifically, we applied profile likelihood analysis (further details can be found in Supplement A.4.4), as it is a widely used non-local frequentist approach to uncertainty quantification in systems biology (Kreutz et al., 2013). The computation of profile likelihoods took seconds, whereas SAEM took on the order of minutes. On the synthetic data, the confidence intervals based on profile likelihoods were comparable to those based on linearization using SAEM for most parameters. Yet, for three variance parameters, the 80% CIs computed with SAEM actually did not cover the true parameter (for all 100 runs of the multi-start), while the CIs computed with profiles from the amortized approach did (Figure 3D).

In conclusion, our amortized approach allows for an efficient and robust uncertainty quantification by computing profile likelihoods. The cheap amortized inference phase is a key advantage, as other frequentist methods do not allow for robust uncertainty analysis due to substantially higher computational costs. Moreover, the efficient evaluation of the population likelihood allows us to perform a full Bayesian analysis as well as we demonstrated in Section 3.4.

#### A.4.4. Profile likelihoods

We show that we can use our approximated population likelihood (5) for uncertainty quantification using the profile likelihood method. To compute confidence intervals from profiles we need to compute the profile likelihood ratio

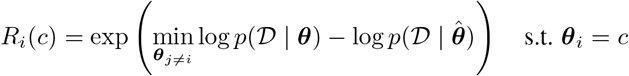

as discussed in (Fröhlich et al., 2014). In our case, we need to compute

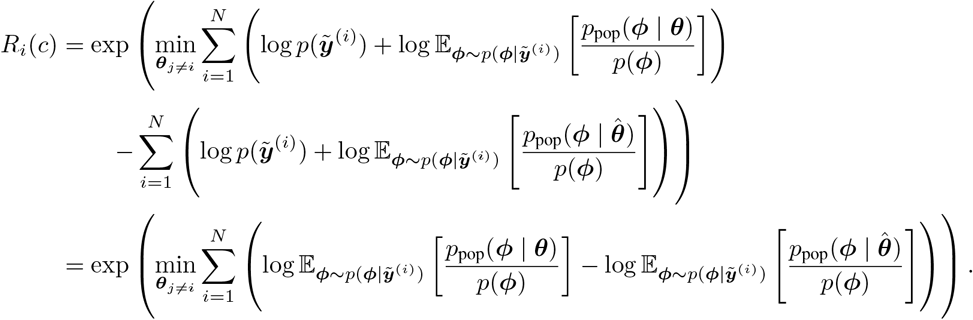

Therefore, we can use the approximation of the population likelihood even though we do not know 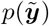. Since the evaluation of this approximation is fast, we can efficiently compute profiles and confidence intervals (see Figure S15). We compute confidence intervals using the implementation in `pyPESTO` (Schälte et al., 2023).

**Figure S15.**
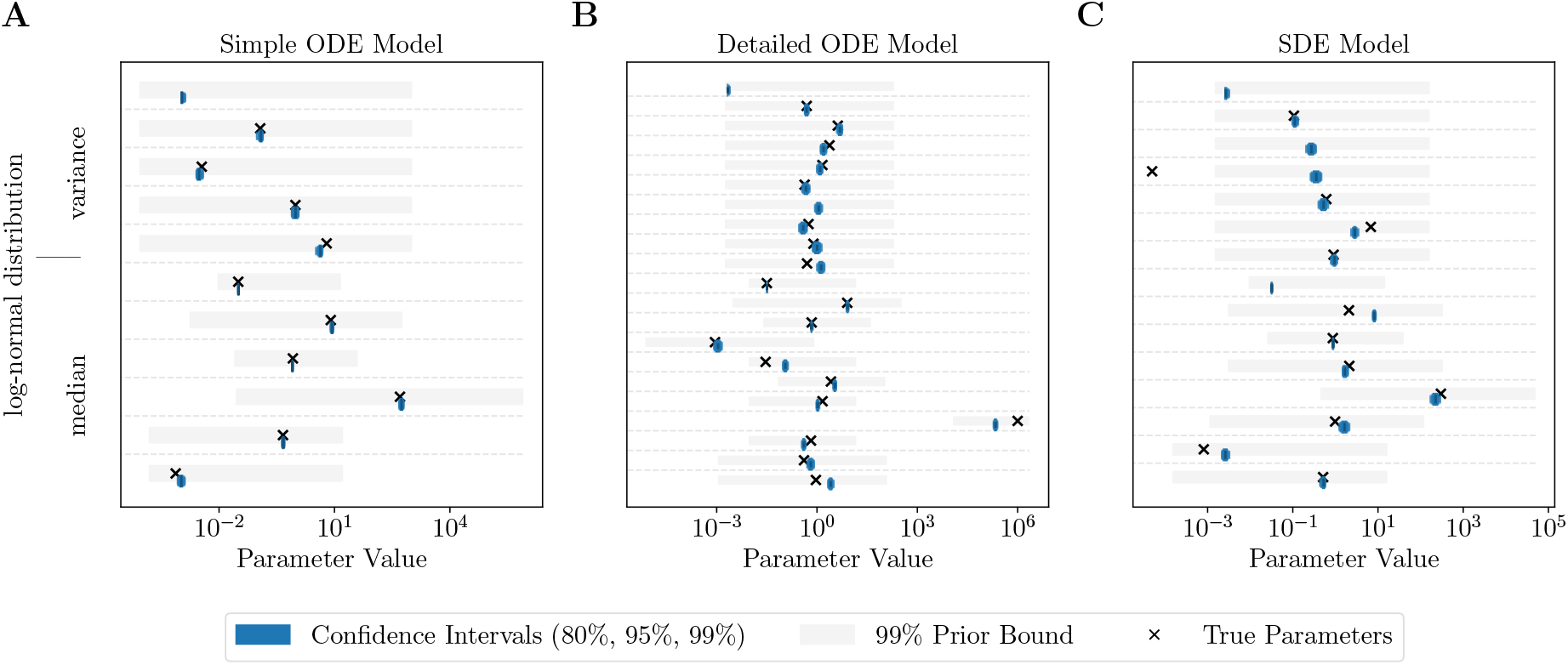
Confidence intervals for the single-cell models on synthetic data. Data was generated by (**A**) the simple ODE model, (**B**) the detailed ODE model and, (**C**) the SDE model. The parameters (median and variance of the log-normal distribution) and CIs (based on profile likelihoods) were then estimated using the amortized approach to NLME models. True parameters, which are 0, are not shown.

#### A.4.5. ODE vs. SDE NLME model

The simple ODE model of the mRNA transfection processes possessed structural non-identifiabilities, meaning that not all the parameters can be determined from the data. Consequently, the ODE model encompasses only the product *k·m*_0_*·scale*, while the SDE model encompasses the individual parameters *k, m*_0_ and *scale*, offering a more detailed representation. Indeed, using our amortizing NLME framework, we were able to identify all parameters of the stochastic NLME model (see Figure S16B).

Further analysis on synthetic data generated by the SDE NLME model showed that the simple ODE NLME model estimated parameters such that the variance of the population was 3 times larger than the true variance, while for the stochastic NLME model the variance is only 1.3 times larger and hence capable of capturing the data more accurately (Figure S17). This, in particular, underlines that a deterministic model can give erroneous results if it inadequately captures the underlying processes.

**Figure S16.**
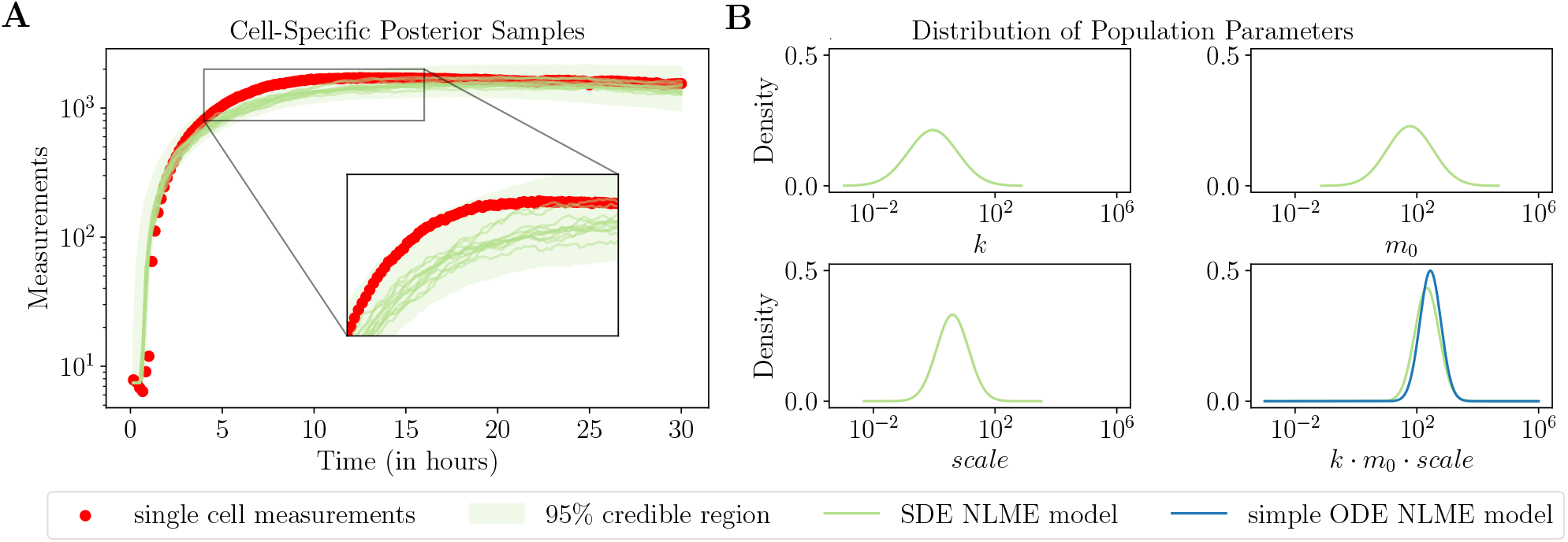
Stochastic NLME model improves identifiability compared to deterministic counterpart. (**A**) Credible regions of a trajectory of the SDE single-cell model estimated by the neural posterior estimator for a real cell. The estimated median of the posterior was simulated 10 times. (**B**) Estimated population distributions for the parameters *k, m*_0_ and *scale* for the SDE NLME model and their product in the simple ODE NLME model.

**Figure S17.**
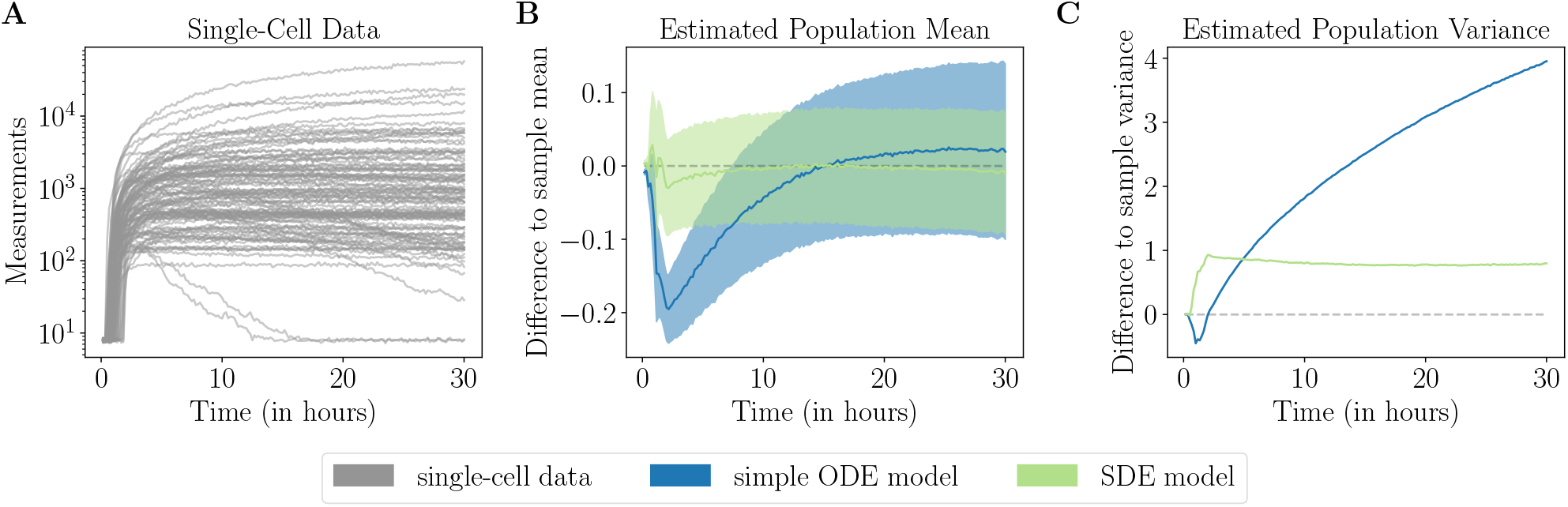
Fit for SDE NLME model on synthetic data. (**A**) Synthetic data describing single-cell translation kinetics after mRNA transfection generated by the SDE NLME model. (**B–C**) Difference of estimated population mean (**B**) and variance (**C**) over time of the SDE and ODE NLME model on synthetic data generated by the SDE model.

### A.5. Specification of the pharmacokinetic model

Over the past two decades, many oral targeted therapies have been developed in the field of oncology, many of which target the angiogenesis of neoplasms, which plays an important role in tumor growth. However, angiogenesis inhibitors generally show high variability between patients, leading to significant differences in exposure (Groenland et al., 2019). Therefore, pharmacokinetic (PK) modeling is required to develop targeted dosing strategies for sub-populations or even in a personalized manner, discover concentration thresholds for toxicity, investigate potential interactions, and guide study planning, among other purposes. Sunitinib, an angiogenesis inhibitor, which belongs to the class of tyrosine kinase inhibitors, was the subject of the population pharmacokinetic model, which is described in more detail below. In the model developed by Diekstra et al., the distribution of sunitinib is described by a single compartment model, while for its metabolite SU12662, a two compartment model was used. Presystemic metabolization in (Diekstra et al., 2017) was described according to the model by Yu et al. by a hypothetical enzyme compartment. The hypothetical compartment was parameterized as follows, with *Q*_*H*_ being the calculated concentration:

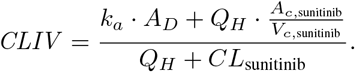

*k*_*a*_ denotes for the absorption rate constant, while *A*_*D*_ and *A*_*c*,sunitinib_ represent the amounts in the dosing or central compartment, respectively. *CL*_sunitinib_ and *V*_*c*,sunitinib_ denote the clearance and volume of distribution of the central compartment of sunitinib in this equation.

The model includes the sex and weight of the patients as covariates. Each patient *i* received a personal medication (*DOS*_*i*_) and was measured over a different period of time and at varying time points. In the following, we present the model for each individual; therefore, the index *i* is removed. The patient’s weight is normalized as follows

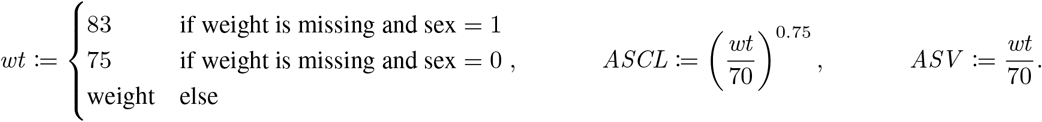

The parameters we want to estimate are 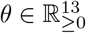, and 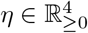, which are incorporated in the ODE model as follows:

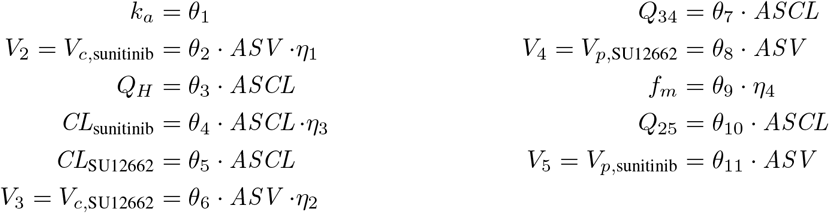

and

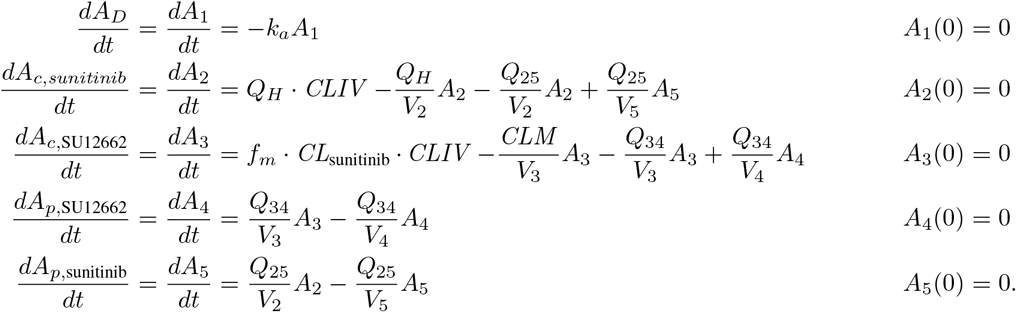

As in the baseline (Diekstra et al., 2017), we fix *θ*_3_ = 80, *θ*_9_ = 0.21, and *θ*_11_ = 588 to get comparable results. Furthermore, whenever a patient takes medication (at 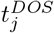), we have

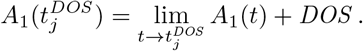

In the noise model we apply a censoring from below by

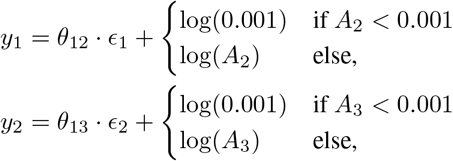

where *ϵ*_1_, *ϵ*_2_∼ 𝒩 (0, *σ*^2^). In (Diekstra et al., 2017), *σ*^2^ = 1 was fixed, therefore we fix it as well. This ODE system is simulated using the Rodas5P solver implemented in the Julia package `DifferentialEquations.jl` (Rackauckas & Nie, 2017).

**Figure S18.**
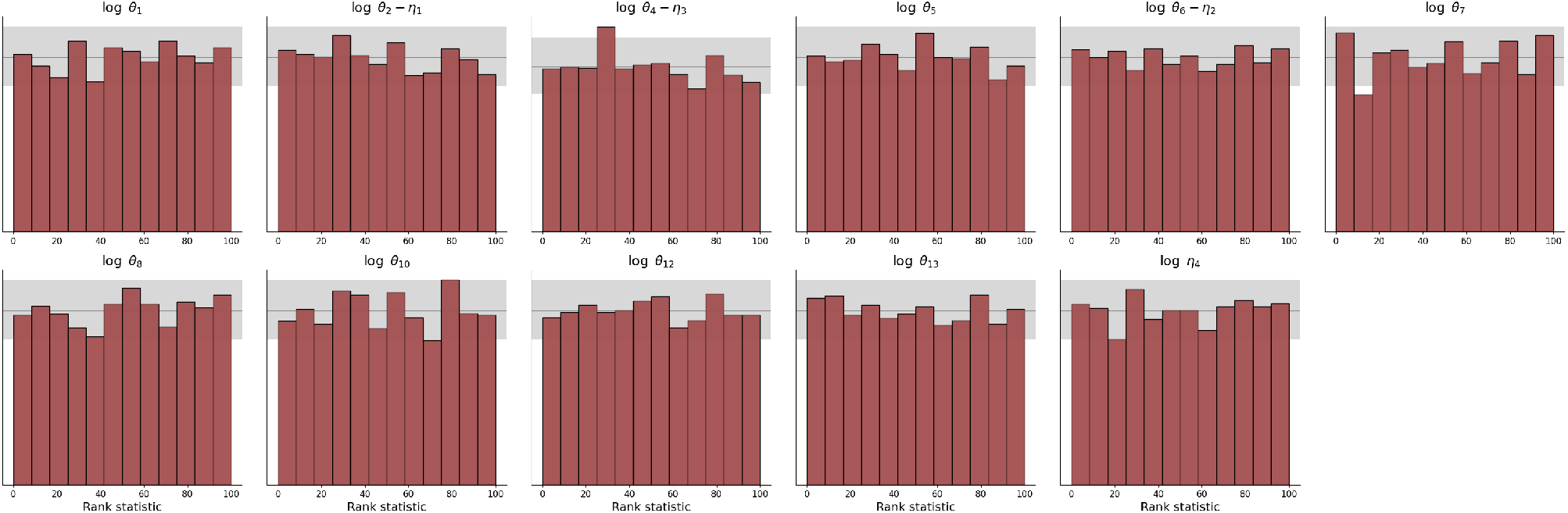
Simulation-based calibration plots of the individual posteriors for the pharmacokinetic model. Incorrect calibration can be seen by deviations from uniformity (bars outside the gray area)

#### A.5.1. Dosing events

In our amortizing framework, covariates such as sex and weight can be treated as part of the population model. If they are instead part of the model ℳ, then they need to be synthetically generated during the simulation phase. This is the case with dosing regimes, which refer to the prescribed schedules and dosages of the medications that are administered to patients. Therefore, we encoded the dosing events as part of the observations, which are given to the summary network together with the simulated measurements. Hence, the observation at each time point *j* consists of a vector (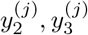, *DOS*_*j*_, *t*_*j*_, *DOS* -*Indicator*_*j*_), where *DOS* -*Indicator*_*j*_ is a binary indicator of a dosing event following the ideas on encoding missing data and time points in (Wang et al., 2023). If a dosing event occurs, the variables are 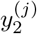 and 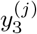 are set to 0, otherwise *DOS*_*j*_ is set to 0. We trained two LSTMs using the split summary network architecture provided in `BayesFlow` (Radev et al., 2023), where each summary network got the input depending on the binary variable. During the simulation phase, we sampled the dosing events and observed time points from the time points and events in the data set because we were only interested in this particular data set. However, one could also generate events and observation time points from a reasonable distribution to be able to amortize over multiple different data sets.

#### A.5.2. Comparison between different estimation methods

We estimated the population parameters using FOCEI, SAEM and our amortized approach. We report the results in Figure S19. Furthermore, we show the measurement trajectories for three different patients to show the underestimation of the population variance of FOCEI (see Figure S20).

Our amortized approach, including all phases, repeating phase (III) 200 times and generating 100, 000 samples from the full population posteriorly, was completed in 27 hours. For an optimization run, we need on average 0.34 minutes. Sampling takes 5.1 minutes.

**Figure S19.**
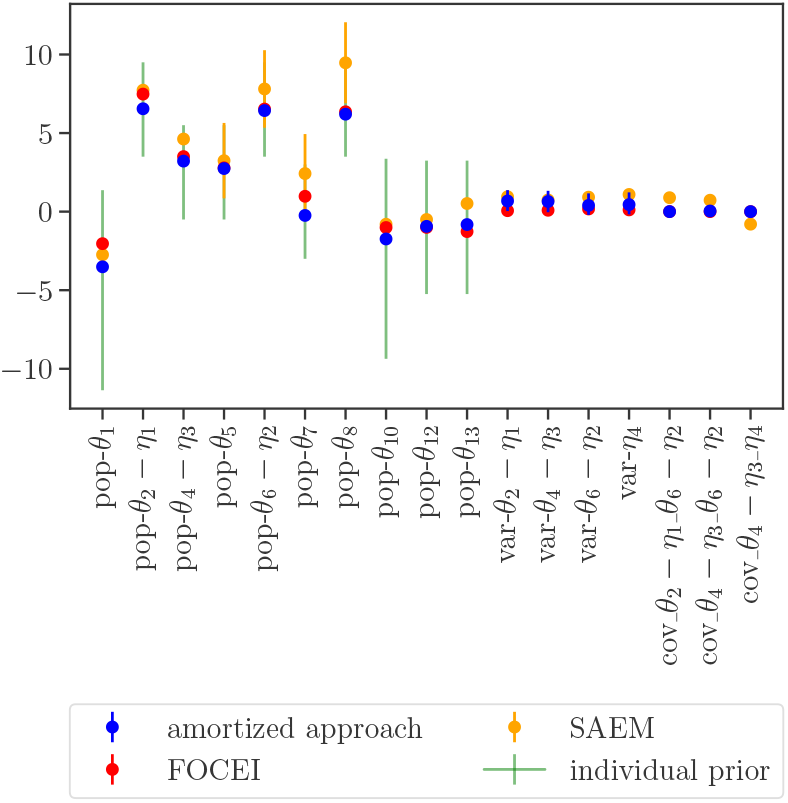
Population parameter estimates for the pharmacokinetic model.

**Figure S20.**
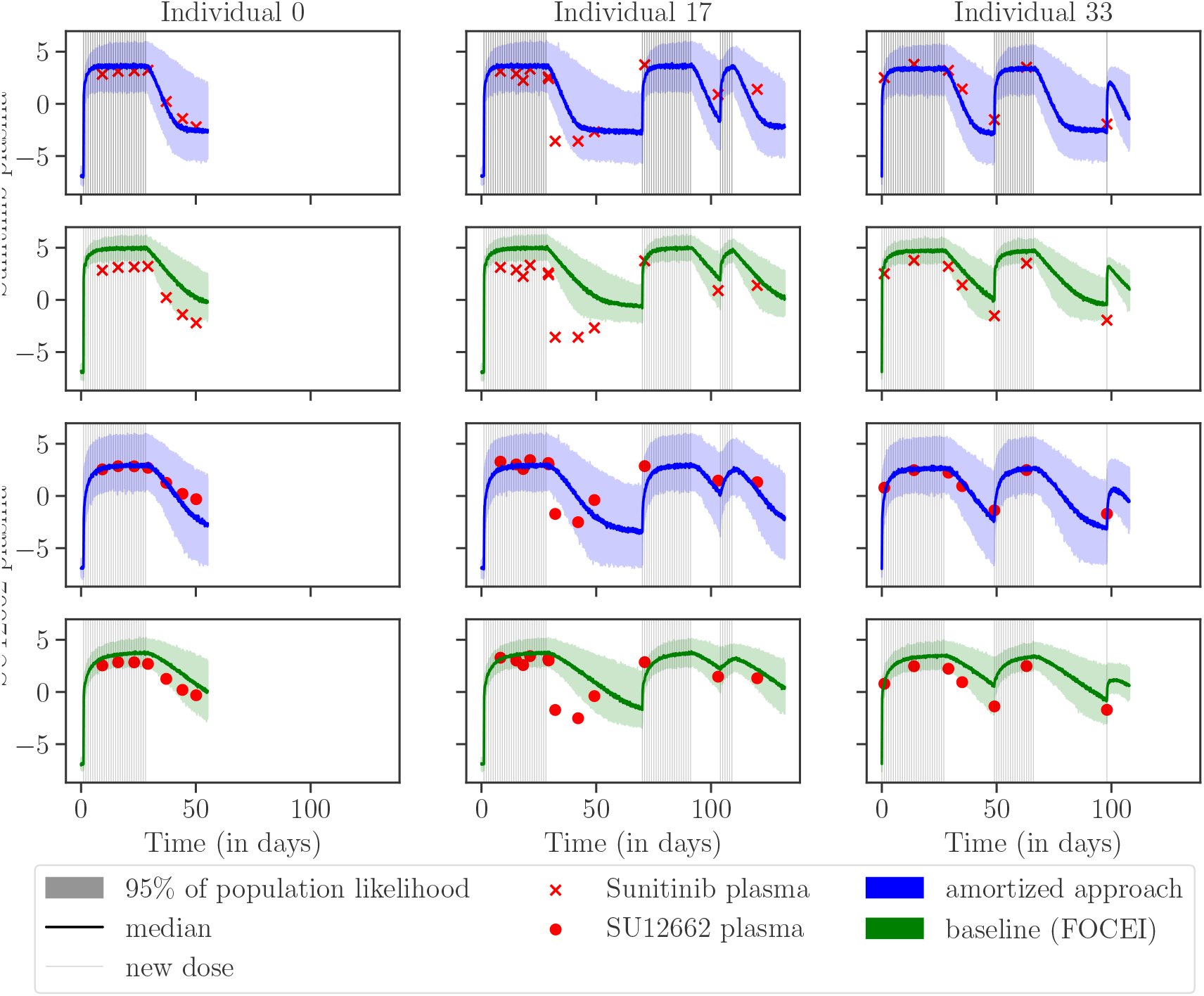
Baseline underestimates variance of measurements. Trajectories of sunitinib plasma and SU12662 plasma measurements for three patients. Simulating samples from the population likelihood convoluted with the noise model using the covariates of this patient based on the estimated parameters of FOCEI and our amortized approach.

## Notes

https://github.com/arrjon/Amortized-NLME-Models/tree/ICML2024

https://zenodo.org/record/8245786

### Competing Interest Statement

The authors have declared no competing interest.

### Summary of Updates

Structure of the paper was improved and a full Bayesian analysis added.

